# Sizes, conformational fluctuations, and SAXS profiles for Intrinsically Disordered Proteins

**DOI:** 10.1101/2023.04.24.538147

**Authors:** Mauro L. Mugnai, Debayan Chakraborty, Abhinaw Kumar, Hung T. Nguyen, Wade Zeno, Jeanne C. Stachowiak, John E. Straub, D. Thirumalai

## Abstract

The preponderance of Intrinsically Disordered Proteins (IDPs) in the eukaryotic proteome, and their ability to interact with each other, folded proteins, RNA, and DNA for functional purposes, have made it important to quantitatively characterize their biophysical properties. Toward this end, we developed the transferable Self-Organized Polymer (SOP-IDP) model to calculate the properties of several IDPs. The values of the radius of gyration (*R*_*g*_) obtained from SOP-IDP simulations are in excellent agreement (correlation coefficient of 0.96) with those estimated from SAXS experiments. For AP180 and Epsin, the predicted values of the hydrodynamic radii (*R*_*h*_s) are in quantitative agreement with those from Fluorescence Correlation Spectroscopy (FCS) experiments. Strikingly, the calculated SAXS spectra for thirty-six IDPs are also nearly superimposable on the experimental profiles. The dependence of *R*_*g*_ and the mean end-to-end distance (*R*_*ee*_) on chain length, *N*, follows Flory’s scaling law, *R*_*α*_ ≈ *a*_*α*_*N* ^0.588^ (*α* = *g*, and *e*), suggesting that globally IDPs behave as synthetic polymers in a good solvent. The values of *a*_*g*_, and *a*_*e*_ are 0.20 nm and 0.48 nm respectively. Surprisingly, finite size corrections to scaling, expected on theoretical grounds, are negligible for *R*_*g*_ and *R*_*ee*_. In contrast, only by accounting for the finite sizes of the IDPs, the dependence of experimentally measurable *R*_*h*_ on *N* can be quantitatively explained using *ν* = 0.588. Although Flory scaling law captures the estimates for *R*_*g*_, *R*_*ee*_, and *R*_*h*_ accurately, the spread of the simulated data around the theoretical curve is suggestive of of sequence-specific features that emerge through a fine-grained analysis of the conformational ensembles using hierarchical clustering. Typically, the ensemble of conformations partitiones into three distinct clusters, having different equilibrium populations and structural properties. Without any further readjustments to the parameters of the SOP-IDP model, we also obtained excellent agreement with paramagnetic relaxation enhancement (PRE) measurements for *α*-synuclein. The transferable SOP-IDP model sets the stage for several applications, including the study of phase separation in IDPs and interactions with nucleic acids.

## 1 Introduction

There are several compelling reasons to quantitatively characterize the biophysical properties of Intrinsically Disordered Proteins (IDPs), which unlike globular proteins, do not have a unique three-dimensional fold, but like homopolymers explore an ensemble of conformations. (1) Despite variations in the precise estimate of the number [1], it is suspected that about half of the human proteins have either a large fraction (> 30%) of disordered residues, or feature long (> 30 or > 40 residues, depending on the location along the chain) disordered regions. [2] (2) IDPs play a central role in a variety of cellular functions because they are not only capable of self-association, but also interacting with multiple partners, including nucleic acids. [3] (3) They organize themselves to form condensates, which are suspected to offer advantages in cellular functions [4–7] such as transcription [8], and enzyme reactions [9, 10]. (4) Although insights into the formation of these biomolecular condensates, in which a high-density droplet coexists with a low-density dispersed phase, may be obtained by borrowing concepts from polymer physics,[11– 13] sequence effects must and do play an important role. For these and other reasons, there is great interest in understanding the physico-chemical properties of IDPs using experiments, simulations, and theories that faithfully take the exact sequence into account.[14, 15]

Despite considerable progress in computer simulations of IDPs [16–24], the need to develop transferable models, with as few parameters as possible, exists. The accuracies of computational models can only be assessed by direct and unbiased comparisons with different experiments. Small-Angle X-ray Scattering (SAXS) profiles, available for a number of IDPs, could be used as the basis for quantifying the accuracy of computational models. There are potentially a large number of energy functions that one could create to describe the shapes and conformational fluctuations of IDPs. Ideally, these models should be transferable, so that reasonably accurate predictions may be obtained for a given sequence and specified external conditions without having to tweak the parameters repeatedly. All-atom simulations, have had some success in describing the sequence-dependent properties of IDPs,[25–28] but their level of accuracy is often dictated by the particular choice of the force-field/water model,[29–33] or even the sampling protocol.[34, 35] In this regard, coarse-grained simulations have emerged as a viable alternative for predicting sequence-dependent properties on large scale. Several coarse-grained models, with different resolutions, have been proposed [19, 22, 36–42]. Recently, we introduced the Self-Organized Polymer (SOP) model for IDPs [43] (abbreviated as SOP-IDP) in which three adjustable parameters in the energy function (see below) were calibrated to reproduce the experimentally determined radii of gyration (*R*_*g*_s) for three IDPs. We then showed that the calculated scattering curves for 24 other IDPs were in good agreement with the experimental SAXS profiles (*I*(*q*)s with *q* being scattering wave vector) without any further fine-tuning of the parameters. However, for some IDPs (Histatin-5 for example), the calculated *I*(*q*) at *q* > 1.7 nm^−1^ (*qR*_*g*_ ≈ 2.29) deviated from experiment (see Fig. S1A in the Supporting Information). Although the calculated and experimental *R*_*g*_s were in good agreement (Fig. S1B in the Supporting Information) with each other, in some instances the predicted *R*_*g*_s were smaller than the experimental values by about 10%. The initial formulation [43] did not include side-chain for alanine, whose inclusion is important, at least for some IDPs. When the alanine side-chain was added to our model, without altering the functional form of the SOP-IDP energy function, we observed significant deviations from experimentally measured *I*(*q*) for α-synuclein. Therefore, we found it necessary to recalibrate the parameters in the SOP-IDP energy function to reduce the deviations from experiments.

Here, we employ a robust optimization procedure to recalibrate the three parameters in the SOP-IDP model [43] using the experimentally determined *R*_*g*_s for five test proteins. Second, using the resulting energy function, we establish using simulations that the *I*(*q*) profiles for 36 IDP sequences are in *quantitative* agreement with SAXS measurements. Third, we show that on average, IDPs behave like random coils, obeying the famous Flory’s scaling law for polymers in good solvent, *R*_*g*_ ∼ *N*^*ν*^ with *ν* ≈ 0.588. Strikingly, the scaling corrections due to the finite sizes of the IDPs are small for *R*_*g*_, and the mean end-to-end distance, *R*_*ee*_. For the hydrodynamic radius, *R*_*h*_, the apparent Flory exponent obtained from simulations is below the value expected for self-avoiding polymers. However, we resolve the incongruity by including finite-size corrections for *R*_*h*_. Sequence effects manifest themselves in the spread of the data around the expected, theoretical curve. Their importance is clearly highlighted if the fine structural details are probed using a hierarchical clustering scheme. In particular, the fractions of compact, semi-compact, and extended conformations vary depending on the sequence and external conditions. Finally, the proposed SOP-IDP model also reproduces the residue-specific paramagnetic relaxation enhancement (PRE) for α-synuclein nearly quantitatively. As testable predictions, we report the I(q)s of AP180 and FUS.

## 2 Computational Methodology

### Energy Function

Following an earlier study [44], and subsequent reports [43, 45–47], each amino acid is represented by two beads. One bead represents the C_*α*_ atom, and the other is located at the center of mass of the side-chain (SC) of the amino acid residues. The only exception is glycine, which is modeled using only one bead located at the backbone. The SOP-IDP interaction potential is,

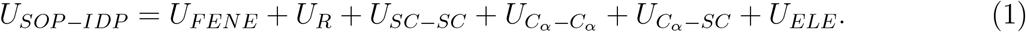

The Finitely Extensible Nonlinear Elastic (FENE) potential (*U*_*FENE*_) accounts for chain connectivity of the backbone C_*α*_ atoms and the chemical bonds between the C_*α*_ and the SC beads. The *U*_*R*_ term is a repulsive potential that prevents unphysical overlap between the beads, acting only between beads that are separated by another bead (1-3 interaction). The terms 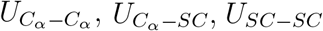 and *U*_*ELE*_ describe the interaction between beads that are separated by at least two other units along the chain (e.g. 1-4, 1-5 … interactions). The Debye-Hückel term, *U*_*ELE*_, accounts for electrostatic interactions between the charged residues. The backbone-backbone (BB), backbone-side-chain (BS), and side-chain-side-chain (SS) interactions are modeled using Lennard-Jones (LJ) potentials,

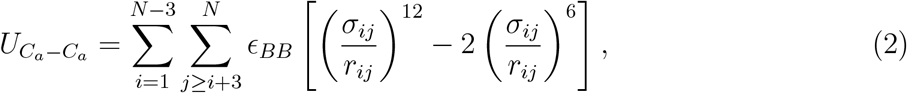

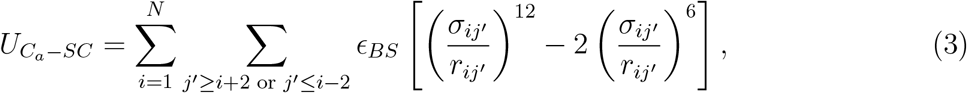

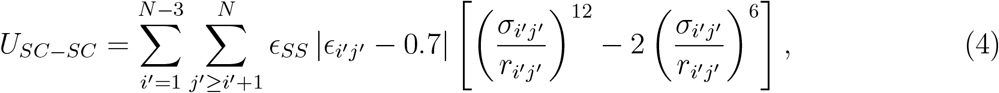

where N is the number of amino acids, indices *i* and *j* refer to residue numbers, where unprimed symbols (*i* and *j*) indicate backbone beads or alpha carbon atoms. The primed counterpart (*i*^*′*^ and *j*^*′*^) indicate SC beads of the corresponding residue, *r*_*ij*_ is the distance between beads *i* and *j*, *σ*_*ij*_ is the sum of the radii of two interacting beads and corresponds to the minimum of the LJ potential. The parameters *ϵ*_*i*′*j*′_ are taken from the statistical potential [48]. There are only three adjustable parameters in the SOP-IDP energy function. They are *ϵ*_*BB*_, *ϵ*_*BS*_, and *ϵ*_*SS*_, which specify the interaction strengths involving the backbone and SCs. We calculate them by minimizing the differences in the calculated and experimental radii of gyration (*R*_*g*_) for a test set of proteins. We then use the optimized values to predict the SAXS profiles for several IDPs, which are not in the traing set.

### Optimization procedure

The values of ϵ_*BB*_, ϵ_*BS*_, and ϵ_*SS*_ are calculated using a minor modification of a parameter optimization (POP) procedure designed to use experimental results to refine force field parameters [49, 50]. Briefly, consider an observable *Ô*_*n*_(Γ), which is a function of the conformational space, Γ, and assume for simplicity that the observable does not depend on the energy of the system. The average of this quantity is,

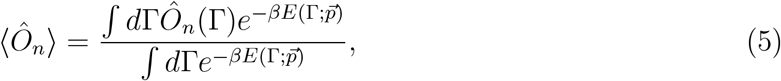

where 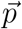 is a set of parameters in the SOP-IDP energy function, 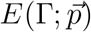 is the energy for the conformation Γ, and *β*^−1^ = *k*_*B*_*T*.

Let *O*_exp,n_ be the experimental value of the observable *Ô*_*n*_(Γ). We compute *R*_*g*_ (which in this case is *O*_exp,n_) for the training set containing five proteins: Histatin-5 (*N*=24), ACTR (*N*=71), Sic-1(*N*=90), K19 (*N*=94), and Osteopontin (*N*=273). The error in estimating a set of *n*_obs_ quantities is calculated using *χ*^2^:

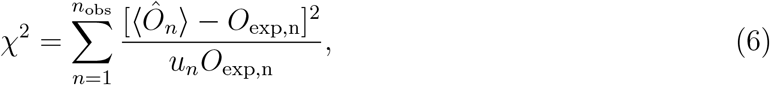

where *u*_*n*_’s are constants that ensure that each element in the sum is dimensionless. Because we use *R*_*g*_ as a target, *u*_*n*_ = 1 nm. The idea is to find a set of parameters 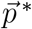 which minimizes χ^2^. We used Newton’s method [51] to obtain the smallest value of χ^2^. This calculation requires the first and second derivatives of the observable with respect to the parameters of the energy function for every step of the optimization procedure. Following the approach outlined previously [49, 50], the derivatives are computed analytically. Their values are obtained by averaging them over a sample of conformations obtained for a given parameter set. After a step, a new set of conformations is sampled with the new parameters, and the procedure is repeated until a satisfying agreement with experiments is achieved. We implemented this procedure with a slight modification. Instead of resampling the conformations after every step (which formally is the correct procedure), we re-weighted the sampled conformations using the new parameters, and took a new step without performing any new simulation. This procedure improves the performance of the algorithm. However, re-weighting becomes inaccurate if the new parameters change significantly from the original values. In order to avoid this scenario, we implemented a confining function that limits the maximum change for each parameter during the re-weighting-based optimization. After a few steps, we re-run the simulations with the new parameter set, making sure that there is only a small difference between the observables obtained via re-weighting the configurations sampled with the previous parameters, and those obtained after sampling with the refined parameters. We repeated this procedure a few times, until convergence was attained. We then used the optimal parameters for simulating the other IDP sequences. The list of IDPs is given in Tables S1 and S2 in the Supporting Information.

#### Simulations

We carried out Langevin dynamics simulations with the SOP-IDP force field using a customized version of the OpenMM code.[52] The equations of motion were integrated using the leap-frog algorithm, using a time-step, *h* = 0.05*τ*. Here, 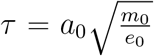 is the natural unit of time, with *m*_0_ = 1 Da, *a*_0_ = 1 ^°^A being the length-scale, and *e*_0_ = 1 kcal/mol being the energy scale. To enhance the sampling efficiency, the simulations were carried out in the underdamped limit, where the viscosity of water was reduced by a factor of 100.[53] For each IDP, simulations were carried out for 2 ×10^9^ steps. The first 3×10^8^ steps were considered as equilibration time, and statistics were collected every 10,000 steps from the rest of the trajectory.

#### SAXS profiles

We calculated the SAXS profiles using all the conformations generated in the simulations using the Debye formula,

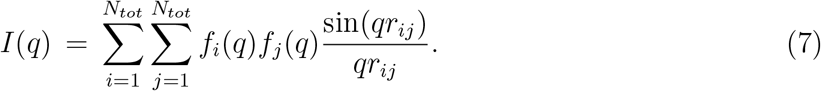

In Eq. 7, *N*_*tot*_ is the total number of interaction sites, and includes both the C_*α*_ and SC beads. The q-dependent form factors, *f*_*i*_(*q*), for the backbone and side-chains are taken from Table S3 of ref [54]. Because Glycine is not explicitly represented as a SC in our model, we calculated the experimental form factor using *f*_*Gly*_(*q*) = *f*_*BB*_(*q*) + *f*_*Gly*−*SC*_(*q*), which operationally means that the distance from another residue to the Glycine BB is equal to the distance from that residue to the Glycine SC. This is not an unreasonable assumption, since the distance between BB and SC within a Glycine residue is ≈1.1 ^°^A (length of a C-H bond for methane, see for instance ref. [55]). It should be noted that not all SAXS experiments report the values of I(0). Moreover, there are considerable uncertainties in extracting I(0) from experiments. Thus, we adjusted *I*(0) to obtain the best agreement between simulation and experiment for *I*(*q*).

#### Radius of Gyration

The radius of gyration, *R*_*g*_, was calculated using:

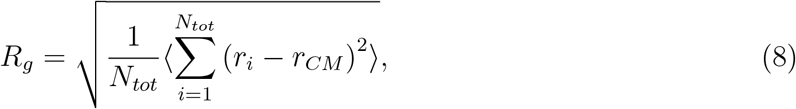

where *r*_*i*_ is the position of bead *i*, and *r*_*CM*_ is the center-of-mass coordinate, and ⟨…⟩ is the ensemble average.

#### Hydrodynamic Radius

We calculated the hydrodynamic radius, *R*_*h*_, using the standard polymer-physics formula,

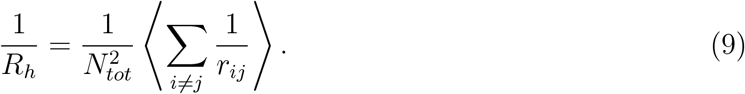

#### Contact maps

Residues *i* and *j* are assumed to be in contact if the distance between their corresponding side-chains is less than or equal to 0.8 nm. Probability contact maps were generated from the time-dependent inter-residue matrices using the following logistic function,

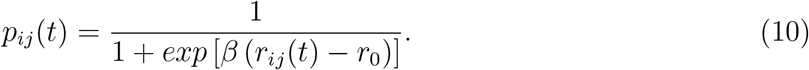

In the above equation, *β* = 500 nm^−1^, and *r*_0_ = 0.8 nm.The ensemble-averaged probability maps, ⟨*p*_*ij*_⟩ are computed by taking an average over all the snapshots *N*_*s*_ in trajectories *N*_*j*_.

### Hierarchical clustering of conformational ensembles

In previous studies [43, 46], we showed that IDP conformational ensembles are highly heterogeneous. Therefore, relying on ensemble-averages alone to ascertain sequence-specific behavior could be misleading, and it does not give a complete picture of the conformational space accessible to IDPs. The representative conformations populating the IDP ensembles were determined using a hierarchical clustering procedure, based on a pairwise distance metric, *D*_*ij*_ = 1 − *χ*_*ij*_, where *χ*_*ij*_, the structural overlap between conformations *i* and *j*, is given by,

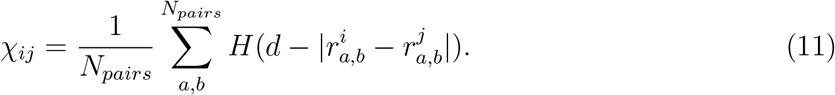

In the above equation, 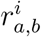 and 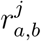 are the distances between the coarse-grained sites *a* and *b* in snapshots *i* and *j*, respectively, and *H* is the Heaviside step function. The tolerance *d*, which accounts for thermal fluctuations, is taken to be 0.2 nm. In Eq. 11, the sum is performed over all the sites that are separated by more than two covalent bonds, and *N*_*pairs*_ is the number of such pairs. The distinct conformational clusters were identified using the Ward variance minimization [56] criterion, as available within the *scipy* module [57].

The contact map for a given cluster, *k* is visualized in terms of the difference map, 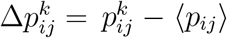, where 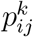 is the probability map for cluster *k*, and ⟨*p*_*ij*_⟩ is the ensemble-averaged probability map. By definition, 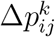 is bound between +1 and –1. Here, we restrict the probability contacts to positive values of 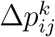 in order to assess how the contact between residues *i* and *j* are enhanced as compared to the ensemble-average.

### Paramagnetic relaxation enhancement

NMR paramagnetic relaxation enhancement (PRE) measurements provide local structural information of the ensemble, which is a consequence of the ⟨*r*^−6^ ⟩ dependence of the PRE between the paramagnetic center and the nucleus of interest, which are usually the backbone amide protons [58]. In the experiments, a paramagnetic probe (such as a nitroxide spin-label) is attached to a mutated cysteine side chain of the protein. The PRE is usually reported as the intensity ratio between two ^15^N-^1^H heteronuclear single quantum coherence (HSQC) spectra in the presence and absence of the probe. Using the conformations generated in the simulations, we calculated the PRE signal from the probe (*p*) to the amino acid (*i*) as follows [32]:

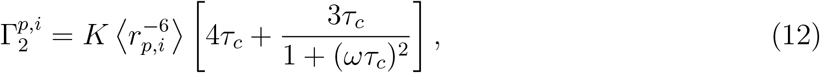

where *K* = 1.23 × 10^−32^ cm^6^s^-2^ is a constant for the nitroxide radical probe, **τ**_*c*_ is the correlation time of the protein reorientation (taken to be 2 ns, which is within the range estimated using atomistic simulations [27]), and *ω*/2π is the Larmor frequency (700 MHz). Because (*ωτ*_*c*_)^2^ ≈ 0.02 is small, it follows from Eq. 12 that 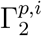 is roughly ∝ *τ*_*c*_, thus making it sensitive to the precise value of **τ**_*c*_.

Due to the coarse-grained nature of the SOP-IDP model, we approximated *r*_*p,i*_ as the distance between the side-chain bead (where the probe attached) to the virtual site that lies between two *i*^th^ and (*i*−1)^th^ C_*α*_ (to mimic the location of the *i*^th^ amide group) (see Fig. 6 of main text). The calculated PRE is converted to the intensity ratio using,

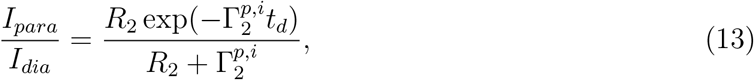

in order to compare directly with experiments. In the above equation, *t*_*d*_ is the total INEPT time of the HSQC experiment (10 ms), and *R*_2_ is the intrinsic transverse relaxation rate estimated from the reduced spectra (4 s^-1^).

## 3 Results and Discussion

### Optimal values of interaction energy scales for side-chains and backbone

The values of *ϵ*_*BB*_, *ϵ*_*BS*_, and *ϵ*_*SS*_ (in unit of kcal·mol^−1^) in Eqs 2-4 after optimizations are:

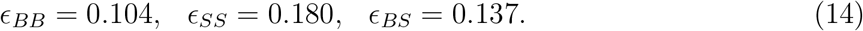

The corresponding values derived previously [43] are (in unit of kcal·mol^−1^) *ϵ*_*BB*_ = 0.12, *ϵ*_*SS*_ = 0.18, and *ϵ*_*BS*_ = 0.24. The largest change is in the value of *ϵ*_*BS*_. Using the new set of parameters greatly improves the agreement of the Histatin-5 SAXS profiles (Fig. S1A, SI Appendix) while maintaining the overall correlation between the calculated and experimental values for the radius of gyration (see below and Fig. S1B).

### IDPs behave as polymers in a good solvent

We first assess the efficacy of the SOP-IDP energy function by comparing *R*_*g*_s with experiments for 36 IDP sequences (see Table S1 in the Supporting Information for a list). The calculated and experimental *R*_*g*_ values are in excellent agreement (Fig. 1A), with the Pearson correlation coefficient being 0.96 (Fig. S1B in the Supporting Information). Based on the results described in Fig. 1B, and Fig. S1e, we surmise that the IDPs behave as homopolymers in a good solvent, that are usually modeled as Self Avoiding Walks (SAWs). (1) The slope of the plot of ln(*R*_*g*_) versus ln(N) (N is the number of residues in a given IDP) is linear with a slope that is consistent with *ν* = 0.588 (Fig. 1B), which is the result obtained for SAWs. Thus, IDPs mimic polymers in a good solvent. To corroborate this observation, we fit to a line *R*_*g*_/*N*^*ν*^ for *ν* in the range 0.45− 0.62. The correct scaling exponent would correspond to the slope that is closest to zero. As shown in Fig S1C, this analysis suggests that the best value of *ν* is in the range 0.60− 0.62. Note that this result is *independent* of the pre-factor, and thus probes only the scaling behavior without the risk of compensation between pre-factor and exponent. Next, by fixing *ν*, we fit the prefactor *a*_*g*_ for *R*_*g*_∼*a*_*g*_*N* ^*ν*^ and obtained *a*_*g*_ ≈ (0.20 0.01) nm. Note that *a*_*g*_ is non-universal, although the extracted value for IDP is close to that found for globular proteins in the unfolded state [63], generated at high denaturant concentrations. (2) In contrast, the same analysis conducted on the hydrodynamic radius, which is measurable using FCS experiments, suggests a scaling *ν* < 0.5 (Fig. S1D). Indeed Fig. 1C shows that *a*_*h*_*N* ^*ν*^ systematically underestimates (overestimates) IDPs with large (small) N. This suggests that finite-size scaling could be significant for the hydrodynamic radius. Accordingly, 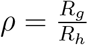 does not match exactly the values expected for a polymer in good solvent (Fig. 2B). (3) The dependence of 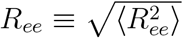 on *N* is also well described by the Flory scaling, with *R*_*ee*_ ≈ *a*_*e*_*N* ^*ν*^, where the extracted parameter is 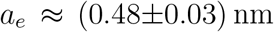 (see Fig S1e). (4) The calculated *R*_*h*_ values for AP180 (N = 593) and Epsin (N=432) agree well with the values obtained using FCS experiments (Table S2, Supporting Information).

**Figure 1:**
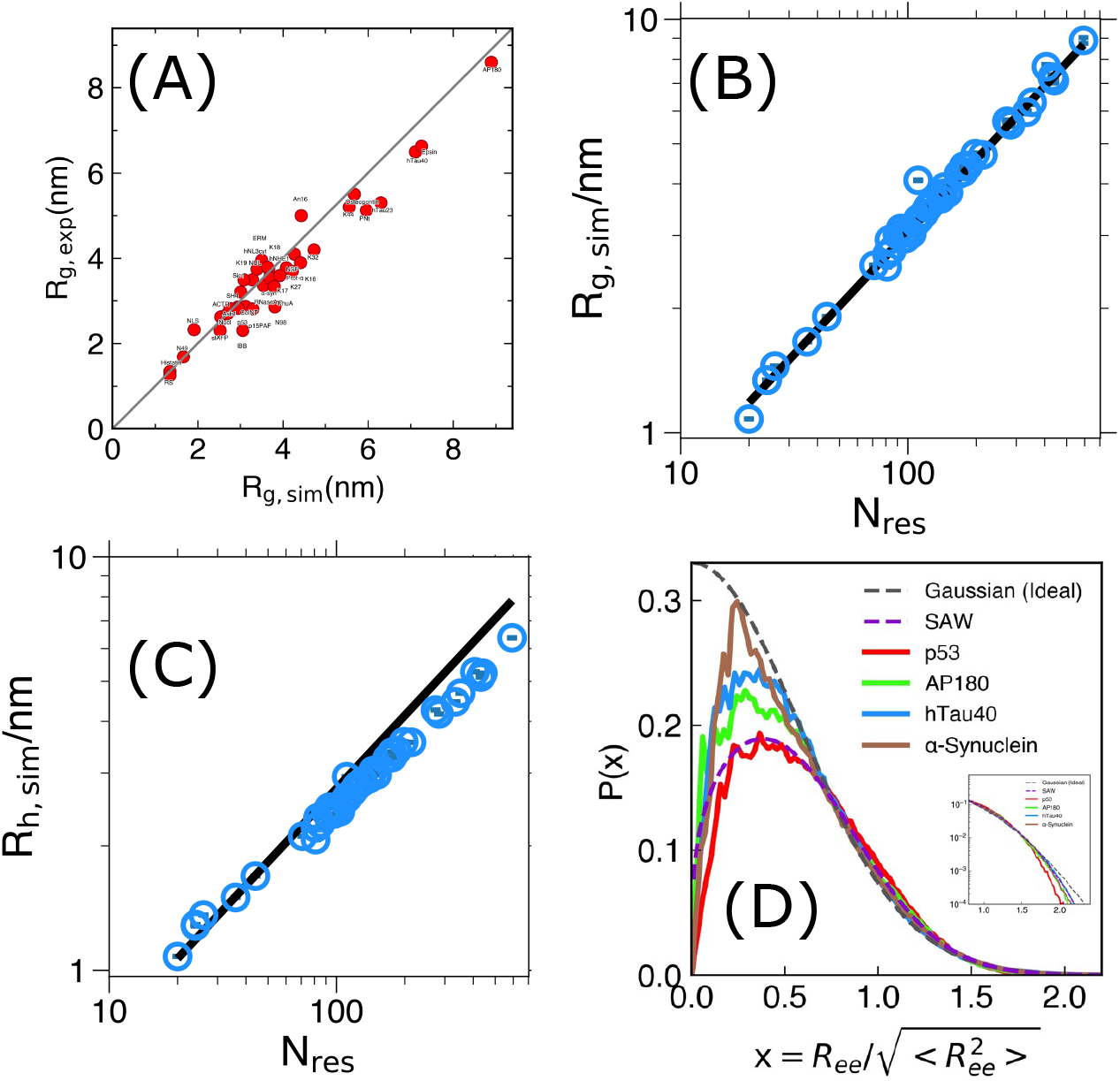
Polymeric features of IDPs: (A) Correlation plot between the radius of gyration calculated from SOP-IDP simulations (*R*_*g,sim*_) and SAXS experiments (*R*_*g,exp*_) for 36 IDP sequences with different lengths and sequence composition. The data are tabulated in Tables S1 and S2 (Supporting Information). The Pearson correlation coefficient is 0.96. (B) Scaling of *R*_*g,sim*_ with the number of residues, *N* as *R*_*g,sim*_ = *a*_*g*_*N* ^*ν*^, with *ν* fixed at 0.588, and *a*_*g*_ = (0.20±0.01) nm. (C) The calculated hydrodynamic radius, *R*_*h*_ compared to Flory’s scaling law with *ν* = 0.588 with *a*_*h*_ = (0.18 ± 0.01) nm. (D) The distributions of the scaled end-end distances for the four IDPs closely resemble the universal theoretical predictions for a self-avoiding random walk (SAW) (Eq.15).

**Figure 2:**
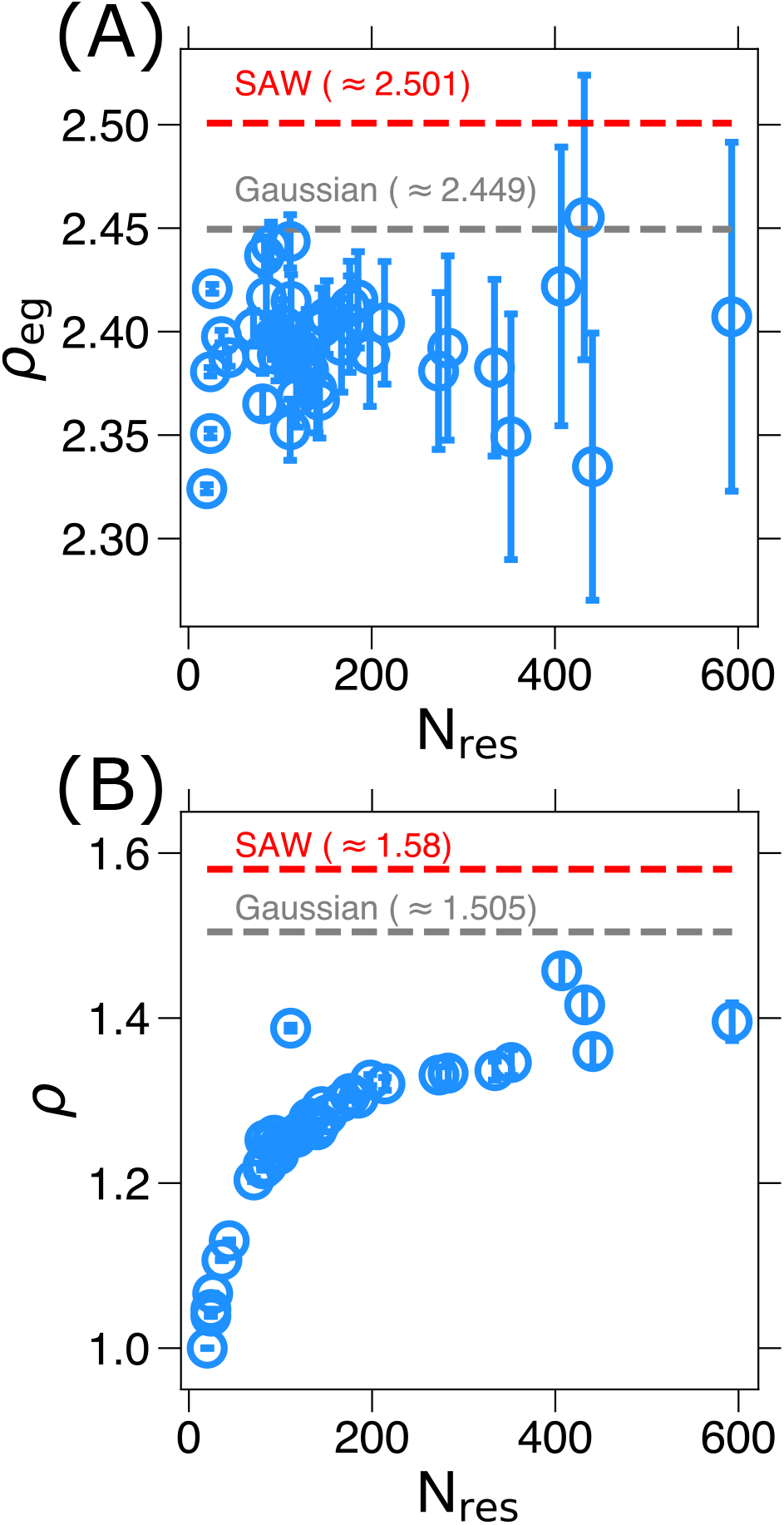
Assessing the universality of *ρ*_*eg*_ and *ρ*: (A) Dependence of *ρ*_*eg*_ = *R*_*ee*_*/R*_*g*_ on *N* calculated using simulations. The values for a Gaussian chain can be computed analytically 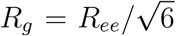 to the leading order, see ref. [59]], so 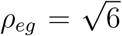). For a SAW the result comes from lattice simulations [60]. (B) Same as (A) except 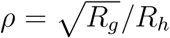 is plotted as a function of N. For a Gaussian chain, analytical calculations [*R*_*h*_ = 3/8(π/6)^1*/*2^*R*_*ee*_ to the leading order, see ref. [61]) give 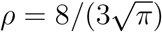, whereas we compare with simulations on a lattice for the SAW [62].

### End-to-end distributions exhibit a universal behavior

If IDPs belong to the SAW universality class, nominally used to characterize homopolymers in a good solvents, then the distribution 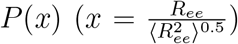 should have a universal behavior [12, 13, 64]. The expression for *P*(*x*) is,

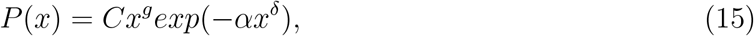

where *C* and α are constants, which are determined using the conditions 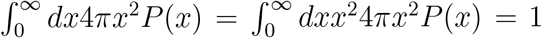. [13] The value of g in Eq. 15, which is zero for a Gaussian chain, is the correlation hole exponent, and 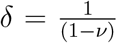. For the four IDPs in Fig. 1D the shapes of the distribution functions *P*(*x*)s resembles the SAW prediction, which is striking given the finite size of the proteins and sequence variations.

### Ratios involving *R*_*g*_, *R*_*h*_, and *R*_*ee*_

For a finite sized SAW, it is well established that there are corrections to scaling for 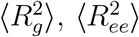 Keeping only the leading order correction in *N*, the scaling correction for *R*_*g*_ may be written as (see for example [62]),

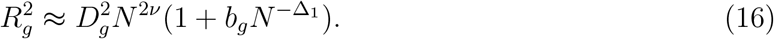

Similarly, 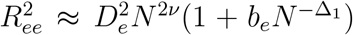 The simplest scaling correction to *R*_*h*_ should have a similar form albeit with an extra term [62],

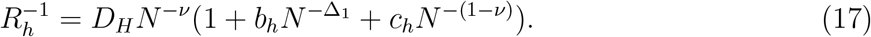

In Eq. 16 and Eq. 17, the coefficients *D*_*g*_, *D*_*e*_, and *D*_*h*_, which are the analogues of *a*_*g*_, *a*_*e*_, and *a*_*h*_, are non-universal, as are the pre-factors *b*_*g*_, *b*_*e*_, *b*_*h*_ and *c*_*h*_ (Eq.16 and Eq. 17). However, for sufficiently large N, the ratios 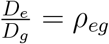 and 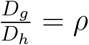 are universal for SAWs [60, 62, 65].

We first calculated *ρ*_*eg*_ and *ρ* for IDPs as a function of *N*. By keeping the corrections to scaling (Eq. 16 and Eq. 17) in mind, we draw the following conclusions from the plots shown in Fig. 2A for *ρ*_*eg*_ and Fig. 2B for *ρ*. (1) Off-lattice simulations for homopolymers for *N*=80 yields *ρ*_*eg*_ = 2.512 [66]. The most accurate estimate for *ρ*_*eg*_, calculated using Monte Carlo simulations for a SAW on a cubic lattice with *N* ≫ 1 (≈ 10^6^), is ≈ 2.501 [62]. The exact result for a Gaussian chain is 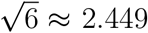 These bounds are reached for the IDPs (Fig. 2A). Instead, the ratio *ρ*_*eg*_ is nearly a constant, in accord with the small correction to scaling obtained for the radius of gyration and the end-to-end distance (see next section). This observation is consistent with our previous work. [43] (2) The dependence of 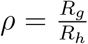 on *N* is relatively smooth. Even for AP180 (*N*=593) the value of *ρ* is less than the exact result for the Gaussian chain (Fig. 2B). The strong dependence on protein size displayed in Fig. 2 suggest that finite size corrections may be relevant, which we examine next. Note that in both cases the theoretical values for the ratios *ρ* and *ρ*_*eg*_ are not reached, which correlates with the observations that, although Flory’s scaling law provides an excellent estimate of the size of the IDPs, ultimately they are heteropolymers. Consequently, the local structural results in subtle deviations from Gaussian or self-avoiding homopolymer description.

#### Corrections to scaling

In order to utilize Eq. 16 and Eq. 17, we set *D*_*x*_ = *a*_*x*_ (*x* =*g*, *e*, or *h*). The values of *a*_*x*_ are obtained using the scaling of *R*_*x*_ as a function of *N* (Fig. 1). Using the values of *a*_*g*_, *a*_*e*_, and *a*_*h*_, and setting Δ_1_ = 0.528 [60, 62], the only free parameter in Eqs. 16,17 are *b*_*g*_, *b*_*e*_, *b*_*h*_ and *c*_*h*_. We determined the first two parameters by fitting the simulation results for *R*_*g*_ and *R*_*e*_ to Eq. 16. The results in Fig. 3A,B and the small values found for *b*_*g*_ and *b*_*e*_ (in fact essentially zero if one accounts for the estimated error bar) show that including finite-size scaling correction has nearly no effect on the quality of the fit.

**Figure 3:**
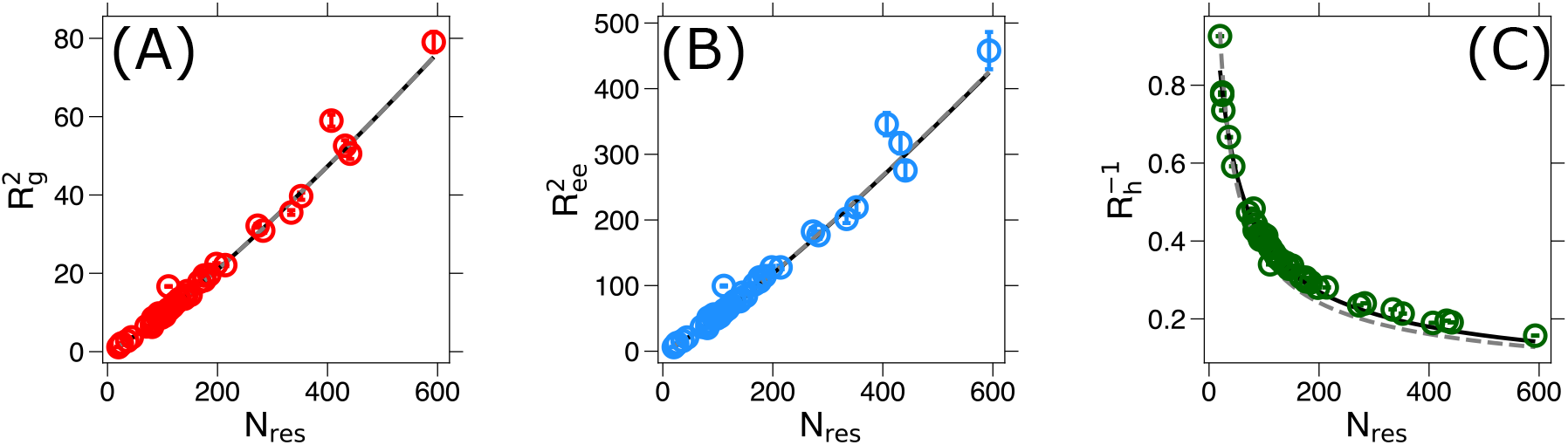
Corrections to scaling for *R*_*g*_, *R*_*e*_, and *R*_*h*_: In all panels, the black lines show the refined fits obtained by including the finite size scaling corrections, whereas the dashed gray lines refer to the same fit shown in Fig. 1. The left and middle panels show the dependence of 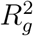 and 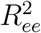 on *N* using Eq. 16. The units for 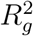 and 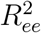 are nm^2^. The best fit values for *b*_*g*_ and *b*_*e*_ are − 0.04 ± 0.45 and − 0.06 ± 0.51, respectively. The one parameter fit gives excellent agreement with experiments. The right panel shows 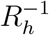 (in unit of nm^−1^) on N. The two parameters in Eq. 17 (*b*_*h*_ = −8.4 ± 6.2 and *c*_*h*_ = 5.6 ± 4.9) were adjusted to reproduce the *R*_*h*_ calculated using simulations. The excellent agreement between simulations and small corrections to scaling further affirms that globally IDPs are similar to polymers in good solvent.

A similar exercise using Eq. 17 using a two parameter fit (*b*_*h*_ and *c*_*h*_) also gives excellent agreement with simulations (Fig. 3C). In this case, the contribution of the correction terms is much larger. Although the error bar is also large, the results suggest that there is a stronger finite-size effect for the hydrodynamic radius. It is worth emphasizing that Eq. 16 and Eq. 17 were obtained for SAWs. That these equations agree almost exactly with the simulation results show that globally the IDP sequences indeed mimic homopolymers in a good solvent.

We should note that corrections to scaling for IDPs are very difficult to measure using experiments. Besides the coefficients *b*_*h*_ and *b*_*e*_ are extremely small. Given the small values of these coefficients, let the relative correction to 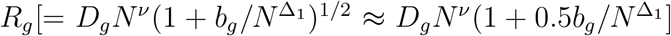 scaling be 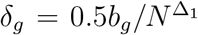. The largest correction is expected for Histatin-5 (N=24). Using we find that *δ*_*g*_ ≈ −3.6 · 10^−3^ %. Thus, for IDPs the Flory scaling *R*_*g*_ is, for all practical purposes very accurate, especially considering the errors in estimating *R*_*g*_ using the Guinier approximation that depends on the range of *q* values used.

### Calculated and experimental SAXS profiles are in good agreement

The calculated SAXS profiles (in the Kratky representation) for six IDPs, and their comparisons to experiments are shown in Fig. 4. The Kratky plots for the other IDP sequences are included in the Supporting Information (Figures.(S2-S6) in the Supporting Information). The SAXS profiles for the six IDPs in Fig. 4 are given in Fig. S7 in the Supporting Information. The calculated Kratky plots for AP180, FUS 1-108, and FUS 1-214 (shown as blue curves in Fig. 4) are predictions whose validation awaits future experiments. The calculated profiles match the experimental results well up to high *q* values for a large numbers of IDPs (Fig. 4 and the Figures (S2-S6) in the Supporting Information), despite substantial variations in sequence, length (N), and experimental conditions. We should emphasize that in these curves we only set an overall scale for the computed scattering profile to set the experimental values. The shape of the curve was not used to train the parameters, nor it was refined using procedures such as ensemble optimization, and thus constitutes a genuine prediction of the model.

**Figure 4:**
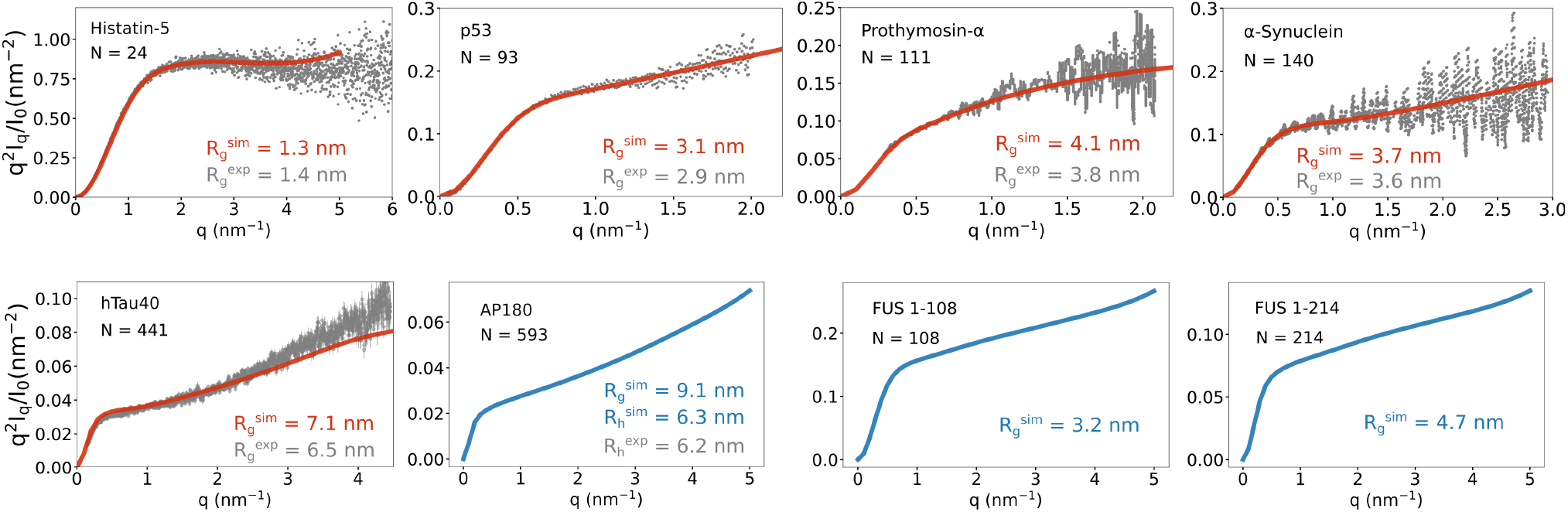
Scattering Curves for IDPs: Small angle X-ray scattering for 6 IDPs in Kratky plot (see Supporting Information for all other IDPs). Comparison between simulated (red) and experimental SAXS (grey) curves. The scattering profiles for AP180, FUS 1-108, and FUS 1-214 (shown in blue) serve as predictions.

### Sequence-specific heterogeneity of IDP ensembles

Although IDPs mimic SAWs closely (Fig. 1, Fig. 2, and Fig. 3), the effect of sequence heterogeneity can be conjectured from the spread of the data. However, a much clearer and more detailed portrayal of sequence effects emerges by examining in detail the ensemble of conformations populated by the IDPs. We visualized the structurally heterogeneous ensemble in the form of a hierarchical partitioning into distinct clusters. In our previous studies, [43, 46] we showed that hierarchical clustering provides a convenient way to glean the finer features of the conformational landscapes that are masked when using ensemble-averages. In Fig. 5, we show the results from the clustering analyses for *α*-synuclein, htau40 and AP180. For each sequence, the minimal partitioning of the conformation space was determined from an elbow analysis of the largest distance jumps in the dendrograms as a function of the number of clusters. Fig. 5 shows the different cluster families are tidily segregated on the two-dimensional landscapes projected onto the *R*_*g*_ and *R*_*ee*_ coordinates, implying that our clustering scheme is robust. In what follows, we provide some additional details related to the partitioning of the conformational space for three IDPs.

**Figure 5:**
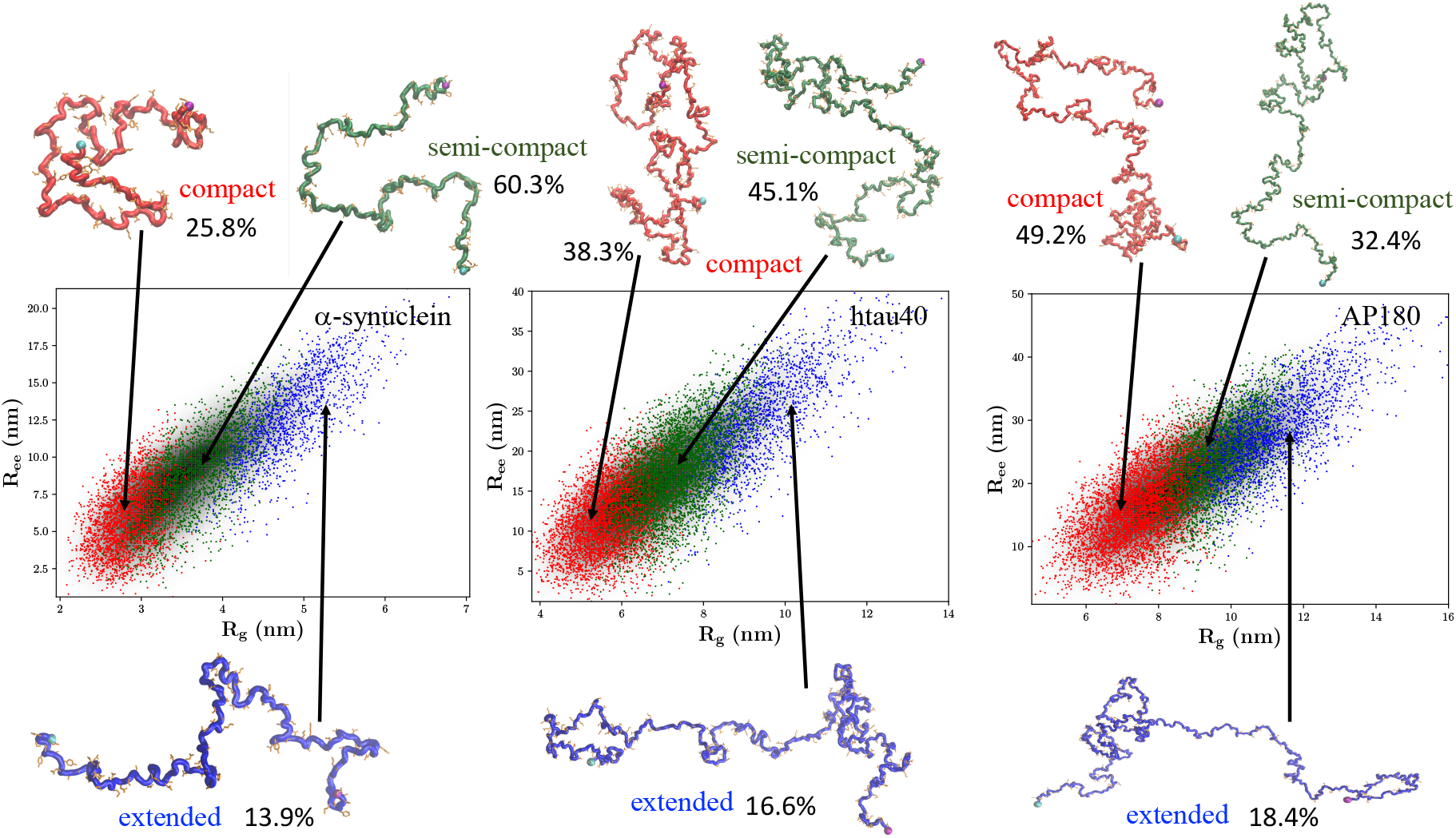
Clustering of conformational ensembles: Hierarchical clustering of the conformational ensembles for *α*-synuclein, htau40 and AP180. For each sequence, the conformational landscape is projected into the *R*_*g*_ and *R*_*ee*_ coordinates. The relative populations of the different clusters, and corresponding snapshots are shown superimposed on the two-dimensional landscapes. The backbones of these representative structures are rendered using different colors. To aid in visualization, the N-terminus residue is marked with a purple sphere, and the C-terminus is denoted as a cyan sphere. The same-color coding is used to distinguish the different clusters on the two-dimensional landscapes.

**Figure 6:**
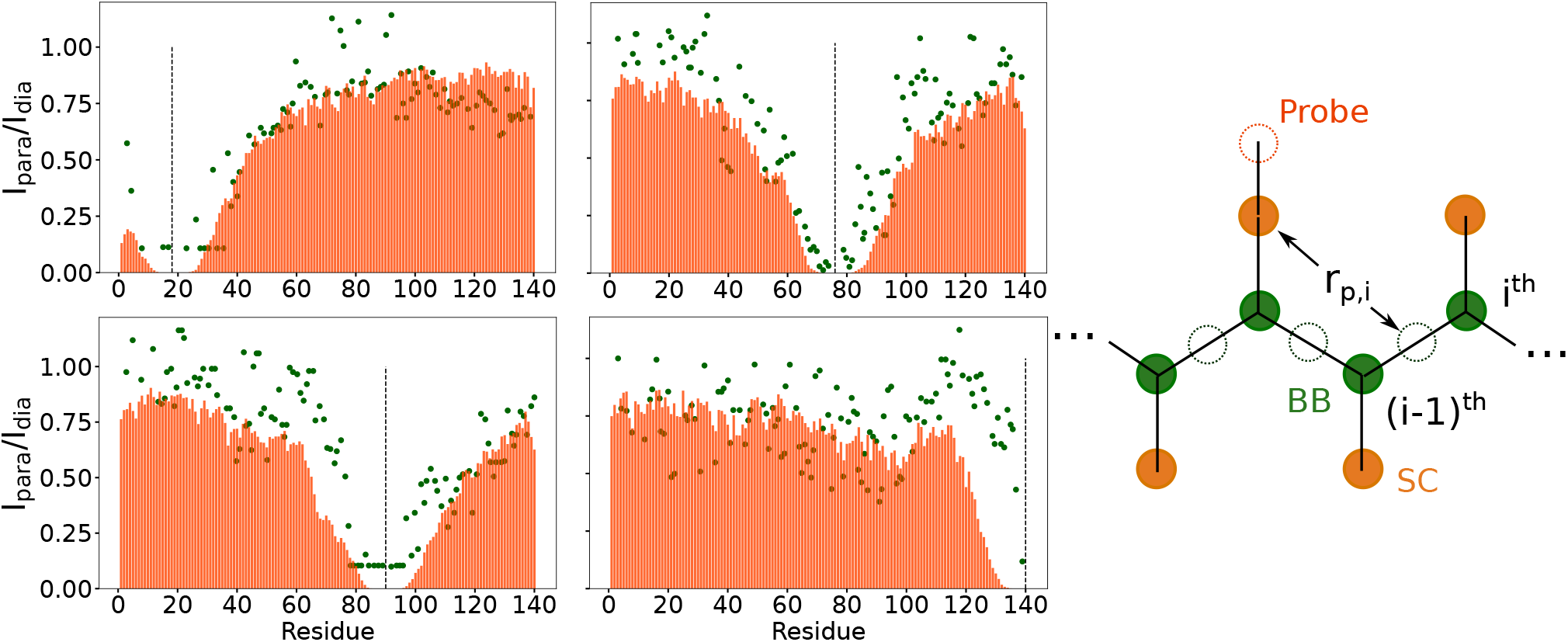
Paramagnetic relaxation enhancement (PRE) of *α*-synuclein. Calculated PREs (orange) are in good agreement with experimental data (green dots) at 4 different locations of the probe (vertical dashed lines) [67, 71, 97]. Here, the distance is calculated from the side chain bead where the probe attached to the imaginary amide groups sits between the backbone C_*α*_ beads.

#### α-synuclein

α-synuclein is a 140-residue IDP, which has been extensively studied using experiments[67–72] and computer simulations, [43, 72–74] because of its key role in synaptic function, as well as its tendency to form aggregates that are implicated in Parkinson’s disease and other synucleinopathies [75]. The α-synuclein sequence can be partitioned into three segments: the N-terminus (residues 1-40); the non-amyloid *β* component (NAC) (residues 41-95); and the C-terminus (residues 96-140). The NAC domain consists of an extended hydrophobic patch, which is a key driver of *α*-synuclein fibrillization. [76, 77]

The ensemble of α-synuclein conformations partition into three clusters, with semi-compact structures having a population of 60.3% (Fig. 5). Compact structures are populated to a lesser extent, and have a population of 25.8%. Fully extended structures are only sparsely populated, and have an occupation probability of 13.9%. Conformations within the compact cluster exhibit different types of residue-residue contacts (Fig. S8A, Supporting Information). In particular, contacts within the NAC domain, which forms the core in α-synuclein fibrils, [78] appear most frequent. Some of the structures within this cluster also show long-range contacts between the N-terminus and the C-terminus, as well as the N-terminus and the NAC domain. Interestingly, contacts between the C-terminus and the NAC domain, which provide a protection mechanism against aggregation, [69, 79] are also evident in some structures within the compact cluster. Many conformations within the semi-compact cluster have weak long-range interactions be-tween the NAC domain and residues 42-55 (often termed as the preNAC domain) within the N-terminus (Fig. S8B, Supporting Information). A recent study by Brockwell and coworkers revealed the presence of such N-terminus-NAC contacts using paramagnetic relaxation enhancement (PRE) measurements, and highlighted their key role in driving α-synuclein aggregation. [80] The contact map corresponding to the most-extended conformational cluster appear largely featureless, although some contacts seem to form transiently within the N-terminus, and the C-terminus and the NAC domains (Fig. S8C, Supporting Information). Our simulations capture the inherent plasticity of α-synuclein, and the partitioning of the conformational space is in accord with previous experimental reports. [67, 72, 81, 82]

#### htau40

The full-length human tau protein (htau40) is a 441 residue IDP. It is localized in the axons of neuronal cells, and regulates the assembly of microtubules.[83] The htau40 sequence can be partitioned into four domains: N-terminus (residues 1-150), proline rich domain (PRD) (residues 151-244), microtubule binding domain (MBD) (245-372), and the C-terminus (residues 372-441). The misfolding of tau and its different isoforms is linked to the formation of neurofibrillary tangles, a hallmark of various taupathies, including Alzheimer’s and other diseases.[84] Because of their ability to undergo phase separation under different conditions, tau and its different truncated variants have been exploited as model systems to understand the regulatory mechanisms underlying the formation of membraneless organelles.[85]. Importantly, recent works suggest that the conformations relevant for phase separation and aggregation may be encoded within the monomer ensemble of tau,[86, 87] much like other amyloidogenic sequences.[46]

Like α-synuclein, there are three major clusters in the conformational ensemble of htau40, corresponding to compact, semi-compact and extended structures (Figure 3). The semi-compact structures dominate the equilibrium with a population of 45.1%. Compact structures are populated to a lesser extent (38.3%), while extended conformations are only sparsely populated (16.6%). Many conformations within the compact cluster exhibit enhanced contacts within the N-terminus, the PRD, and the MBD (Supporting Information, Figure S9A). The sequence of htau40 switches from predominantly acidic to mostly basic around residue 118. [88] This suggests that attractive electrostatic interactions might lead to local compaction of the chain. Accordingly, we observe a square-like pattern in the contact map around residues 100-150, which is clear in the compact ensemble (Supporting Information, Fig. S9A), visible in the semi-compact cluster (Supporting Information, Fig. S9B), and nearly absent for the more expanded ensemble (Supporting Information, Fig. S9C). Long-range contacts between the N-terminus and the PRD, which are key mediators of tau phase separation,[88, 89] exist in only a few conformations within the compact cluster (Supporting Information, Fig. S9A). Conformations within the most populated semi-compact cluster are devoid of any long-range contacts, and only local ordering is visible within the N-terminus, the PRD, and the MBD in some of the structures (Supporting Information, Fig. S9B). Fully extended conformations, which are only sparsely populated within the tau monomer ensemble, do not exhibit any residue-residue contacts (Supporting Information, Fig. S9C).

#### AP180

The endocytic protein AP180 plays an important role in clathrin assembly.[90, 91] AP180 is an adaptor protein of the clathrin-mediated endocytic pathway. Expressed primarily in neuronal cells, AP180 is responsible for linking phosphoinositide lipids to the clathrin coat and for recruiting SNAREs, the transmembrane proteins responsible for synaptic vesicle fusion, into clathrin-coated vesicles. [92] AP180 consists of a structured N-terminal ANTH (AP180 N-terminal Homology) domain, followed by an intrinsically disordered C-terminal domain of approximately 571 amino acids (Rattus norvegicus) [93]. The C-terminal domain of AP180 has been used as a model system to understand how IDPs induce membrane curvature.[94, 95] The ANTH domain is responsible for recruiting phosphoinositide’s and SNAREs, while the C-terminal domain participates in a network of weak, multi-valent contacts with clathrin and other components of the endocytic machinery [96]. The C-terminal domain has a moderately negative net charge per residue of -0.06, where the majority of charged residues are concentrated in the first third of the domain, which has a more significant net negative charge per residue of -0.11 [93].

We simulated a 593-residue fragment, consisting of the AP180 C-terminal domain (residues 23-593) with a HIS-tag linker (residues 1-22). The calculated value of the hydrodynamic radius (6.37 nm) is in near quantitative agreement with the estimate (6.0 nm) from fluorescence correlation spectroscopy experiments (Table S2 in the Supporting Information).[93] The *R*_*g*_ values from simulations and the Flory scaling relation are 8.89 nm and 8.15 nm, respectively. These specific comparisons suggest that AP180, like other IDPs simulated here, behave as homopolymers in good solvent.

Hierarchical clustering reveals that, unlike *α*-synuclein and htau40, the conformational ensemble of AP180 is dominated by compact structures, having an overall population of ≈ 49%. Semi-compact structures, which dominate the equilibrium population of both α-synuclein and htau40, are sampled less frequently for AP180 (population≈32.4%). Extended conformations are only sparsely population (≈18.4%). The long-range contacts between the different residues are not readily discernible from the contact maps of these three clusters (Supporting Information, Fig. S10), which result from a minimal partitioning of the conformational space. To gauge key structural details, we also performed clustering at a finer resolution. This divides the three major clusters (compact, semi-compact and extended) into additional subfamilies. We find that most structures within the compact cluster exhibit various types of long-range contacts, with a majority of them involving residues near the C-terminus (Supporting Information, Fig. S11A). The semi-compact cluster also consists of many conformations displaying long-range residue-residue contacts. However, these are mostly localized in the central segment (≈residues 210-270) of the peptide (Supporting Information, Fig. S11B). The conformations within the extended cluster are mostly RC-like, resulting in a featureless contact map (SI Appendix, Fig. S11C), but some of the subfamilies consist of structures exhibiting contacts between residues near the C-terminus (Supporting Information, Fig. S11D).

Overall, the results described in Fig. 5 illustrate that there are important details in the conformational ensemble of the ground states of IDPs that are masked in the global properties. To ensure that the hierarchical clustering scheme unambiguously reflects the sequence-specific features of IDPs, we simulated a SAW with *N* = 140 (see Supporting Information for details). The SAW ensemble appears more homogeneous, and is segregated into two classes (compact and extended) with nearly equal equilibrium populations (Supporting Information, Fig. S12). Therefore, the more nuanced partitioning of the conformational space displayed in Fig. 5 is a reflection of the sequence.

### Paramagnetic relaxation enhancement (PRE) shows great recovery of local structures

In order to assess the accuracy of the model in predicting the local structures, we compared the calculated PRE with experimental measurements. PRE is an excellent tool to provide local structural information because of the long range (⟨*r*^−6^⟩) nature of the PRE effect, allowing for the detection of transient interactions up to 35 ^°^A[58]. We choose α-synuclein to calculate PRE because of the wealth of available experimental and simulated data, which can be directly compared with our simulations.[27, 32, 67, 71, 97, 98]. Fig. 6 shows the intensity ratios of PRE effects for α-synuclein where the nitroxide probe is located at 4 different positions. The agreement between the calculated and experimental PRE is good. This suggests that the local structural features in our simulations reflect the experimental structural ensemble of α-synuclein.

## Conclusions

Simulations using a transferable coarse-grained SOP-IDP model show that the SAXS profiles for thirty six IDPs are accurately reproduced. The calculated hydrodynamic radii for AP180, the largest IDP simulated here, and Epsin (N=432) agree quantitatively with FCS experiments (relative errors of about 6% and 9%, respectively). In addition, the agreement between simulations and paramagnetic relaxation enhancement measurements is excellent. With these achievements as a background, we summarize the lessons that are likely to be widely applicable for other IDPs. (i) The global properties, such as the radius of gyration and the mean end-to-end distance as a function of the number of residues obey the Flory scaling law accurately. As a result, finite-size scaling corrections are small, even for the 25-residue Histatin-5, in particular for the radius of gyration and the end-to-end distance. However, for the hydrodynamic radius finite-size corrections are required to improve the agreement with Flory’s scaling law. Thus, the Flory scaling law suffices to estimate the sizes of IDPs without the need for experiments. (ii) The spread of the data around the theoretical curve signals that sequence-specific features provide small but measurable deviations around the expected IDP size. In addition, universal ratios between the radii of gyration and end-to-end distances, or between the radii of gyration and the hydrodynamic radii do not converge to the theoreticallay values expected for homopolymers. (iii) A fine-grained analysis further highlights and explains the sequence-dependent properties of IDPs. For instance, by probing the conformational ensemble for α-synuclein, htau40, and AP180 using hierarchical clustering, we found that the structure space partitions into clusters that have distinct properties. The extent of compaction between the clusters, and their weights are sequence dependent. It is likely that the populations of a specific structure are more likely to be functionally relevant, as has been shown in the aggregation of Aβ peptides [46] and Fused in Sarcoma [45]. (iv) Without tweaking any of the parameters, our simulations reproduce, semi quantitatively, experimental paramagnetic relaxation measurement at four probe locations in α-synuclein. All these results show that the SOP-IDP model is transferable.

### Comparison with other models

One can construct a large number of energy functions that can reproduce chosen experimental data. However, the space of energy function would likely narrow if the efficacy is assessed by comparison with multiple measurements. With the caveat that all empirical energy functions are arbitrary, we compare the SOP-IDP model with other models, which have been used to determine single-chain and multi-chain properties. The recent models [19, 24, 40, 41] use the single bead (referred to as C_*α*_ carbon model in protein folding) to represent amino acids. Because the *R*_*g*_ obtained in the initial version [19] produced compact structures, which disagreed with experiments, the stickiness parameter was tuned to obtain better agreement with *R*_*g*_ and PRE measurements [40]. In the Mpipi [41] model, which is also a single bead model, the interactions between the beads were constructed by matching to atomically detailed simulations. The calculated 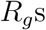 for 17 IDPs agreed well (correlation coefficient of 0.90) with experiments. In addition, the computed coexistence curves, as a function of temperature and concentration, for IDPs were in agreement with the available experiments. The SAXS profiles were not reported in previous studies. Given our finding that *R*_*g*_ is accurately predicted using the Flory scaling relation, even without finite size corrections, more stringent tests have to be performed to assess the validity of these models for single chain properties. At the very least, ii would be important to compare SAXS profiles for several IDPs. Previous studies have shown that [29–33] the *R*_*g*_ estimates from simulations are sensitive to the details of the force field, with the error relative to experimental values being approximately 50% for some sequences (for e.g. Sic1 and α-synuclein modelled with the CHARMM36m force field, Table S3, Supporting Information). Reparametrization of the force field parameters [33] does rectify the over-compaction problem in certain cases. The SOP-IDP simulations fare better across the board (Table S3, Supporting Information). In Table S4 (Supporting Information), we show a systematic comparison of the *R*_*g*_ values obtained for α-synuclein, with different force fields and water models. Most all-atom force fields underestimate the *R*_*g*_ of α-synuclein, although reweighting the ensemble, which is erroneous, improves the agreement with experiment [27]. On the other hand, SOP-IDP accurately describes the size of α-synuclein without requiring any reweighting of the conformations. Its performance is comparable to AMBER19SB/TIP4PD,[99] and the bead-necklace model,[100] which provide accurate estimates of α-synyuclein’s *R*_*g*_.

The SOP-IDP model differs in a few respects from other coarse-grained models. (1) Each amino acid uses two beads (*C*_*α*_-SC in the protein folding context [101]). In contrast to the coarse-grained models [24, 40, 41], which have many parameters, the SOP-IDP has only three parameters whose values were determined by fitting the calculated values to the measured *R*_*g*_ for five proteins in the training set. (2) We simultaneously obtained agreement with experiments for *R*_*g*_, *I*(*q*), and PRE. None of the models have compared *I*(*q*) with experiments. (3) The use of two beads, one for *C*_*α*_ and the other for side chains, is likely to better represent the properties of both single chains and condensates than one bead representation.

## Supporting Information

A list of all IDP sequences considered in this work, along with the *R*_*g*_ values obtained from simulations, and those estimated from SAXS experiments; comparison of hydrodynamic radii, *R*_*h*_ from simulations, and those estimated from FCS measurements; comparison of the *R*_*g*_ estimates and SAXS profiles from the current and earlier force-field parameters; comparison of the *R*_*g*_ values obtained from SOP-IDP simulations, and those from all-atom simulations, for different IDP sequences; *R*_*g*_ values of α-synuclein predicted using different force-fields; SAXS profiles of IDPs in the Kratky representation; scattering profiles of Histatin-5, p53, Prothymosin-*α*, α-synuclein, htau40 and FUS 1-214; dendrograms displaying the hierarchical organization of clusters in the *α*-synuclein and its corresponding SAW ensemble; residue-residue contact maps for the different conformational clusters of α-synuclein, htau40 and AP180.

Scripts and relevant input files that were used to generate the data, as well as detailed instructions regarding the implementation of the SOP-IDP force-field within OpenMM, are available from https://github.com/tienhungf91/SOP_IDP2.

## Acknowledgement

We are grateful to Dr. Farkhad Maksudov for useful comments. We acknowledge the Texas Advanced Computing Center (TACC) for providing the necessary computing resources. This work was supported by a grant from the National Institutes of Health (GM-107703), the National Science Foundation (CHE 2320256), and a grant from the Welch Foundation (F-0019) administered through the Collie-Welch Regents Chair.

The authors declare no competing interest.

