## Supporting Information for "Sizes, conformational fluctuations, and SAXS profiles for Intrinsically Disordered Proteins"

Table S1: List of Intrinsically disordered proteins studied in this work.
† In ref[1], the value of $R_g$ estimated from the distance distribution $P(r)$ distribution is 1.38 nm, and from a Guinier fit to the SAXS profile is 1.33 nm. To compare with our simulations, we took an average of these two values.

| Protein | Length | Salt conc. (mM) | Temp (C) | $R_{g(sim)}(nm)$ | $R_{g(exp)}(nm)$ | Reference |
|---|---|---|---|---|---|---|
| GS | 20 | 150 | 25 | 1.08 | N/A | N/A |
| Histatin-5 | 24 | 150 | 25 | 1.34 | 1.35† | [1] |
| RS repeat | 24 | 150 | 25 | 1.34 | 1.26 | [2] |
| HIV | 26 | 150 | 25 | 1.45 | N/A | N/A |
| N49 | 36 | 150 | 25 | 1.66 | 1.69 | [3] |
| NLS | 44 | 150 | 25 | 1.91 | 2.33 | [3] |
| ACTR | 71 | 200 | 5 | 2.54 | 2.63 | [4] |
| sfAFP | 81 | 150 | 25 | 2.52 | 2.31 | [5] |
| Nucleoporin | 81 | 150 | 25 | 2.70 | 2.70 | [3] |
| Ash1 | 83 | 150 | 25 | 2.93 | 2.85 | [6] |
| SH4 | 85 | 150 | 25 | 2.76 | 2.82 | [7] |
| Sic1 | 90 | 150 | 25 | 3.01 | 3.21 | [8] |
| p53 | 93 | 150 | 25 | 3.12 | 2.87 | [9] |
| IBB | 97 | 150 | 25 | 3.06 | 3.12 | [3] |
| ColNT | 98 | 400 | 4 | 2.95 | 2.83 | [10] |
| Prothymosin-$\alpha$ | 111 | 150 | 25 | 4.08 | 3.78 | [11] |
| p15PAF | 111 | 150 | 25 | 3.29 | 2.81 | [12] |
| NUL | 112 | 150 | 25 | 3.28 | 3.50 | [3] |
| hNL3cyt | 118 | 250 | 20 | 3.39 | 3.00 | [13] |
| ERM | 122 | 150 | 25 | 3.50 | 3.96 | [14] |
| RNaseA | 124 | 150 | 25 | 3.54 | 3.30 | [15] |
| hNHE1 | 131 | 200 | 25 | 3.67 | 3.75 | [4] |
| $\alpha$-Synuclein | 140 | 200 | 25 | 3.68 | 3.55 | [16] |
| FhuA | 143 | 150 | 25 | 3.79 | 3.34 | [15] |
| N98 | 151 | 150 | 25 | 3.81 | 2.86 | [3] |
| NSP | 176 | 150 | 25 | 4.27 | 4.10 | [3] |
| An16 | 185 | 100 | 25 | 4.43 | 5.00 | [17] |
| Osteopontin | 273 | 80 | 25 | 5.67 | 5.50 | [18] |
| K44 | 283 | 150 | 15 | 5.56 | 5.20 | [19] |
| PNt | 334 | 150 | 25 | 5.96 | 5.13 | [15] |
| K19 | 99 | 150 | 15 | 3.08 | 3.50 | [19] |
| K18 | 130 | 150 | 15 | 3.63 | 3.80 | [19] |
| K17 | 145 | 150 | 15 | 3.93 | 3.60 | [19] |
| K27 | 167 | 150 | 15 | 4.22 | 3.70 | [19] |
| K16 | 176 | 150 | 15 | 4.41 | 3.90 | [19] |
| K32 | 198 | 150 | 15 | 4.73 | 4.20 | [19] |
| hTau23 | 352 | 150 | 15 | 6.30 | 5.30 | [19] |
| hTau40 | 441 | 150 | 15 | 7.11 | 6.50 | [19] |
| FUS | 214 | 150 | 25 | 4.70 | N/A | N/A |
| FUS 1-108 | 108 | 150 | 25 | 3.21 | N/A | N/A |
| FUS 111-214 | 104 | 150 | 25 | 3.03 | N/A | N/A |

Table S2: Comparison of Hydrodynamic Radius, $R_h$, from SOP-IDP simulations and from FCS measurements for selected IDP sequences.

| Protein | Length | Salt conc. (mM) | Temp (C) | $R_{g(sim)}(nm)$ | $R_{h(sim)}(nm)$ | $R_{h(exp)}(nm)$ | Ref |
|---|---|---|---|---|---|---|---|
| AP180 | 593 | 150 | 25 | 8.89 | 6.37 | 6.0 | [20] |
| Epsin | 432 | 150 | 25 | 7.25 | 5.12 | 4.7 | [21] |
| Amphiphysin | 407 | 150 | 25 | 7.68 | 5.27 | | [22] |
| Amphiphysin | 382 | 150 | 25 | | | 5.3 | [22] |

| Protein | Experiment (nm) | SOP-IDP (nm) | AMBER99DISP (nm) | CHARMM36m |
|---|---|---|---|---|
| ACTR | 2.63 | 2.54 | 2.12 | 1.86 |
| $\alpha$-synuclein | 3.50 | 3.67 | 3.67 | 1.84 |
| Sic1 | 3.21 | 3.03 | 2.44 | 1.49 |
| Ash1 | 2.85 | 2.94 | 3.16 | 2.34 |

Table S3: Comparison of the $R_g$ values for different IDP sequences obtained with SOP-IDP, and the all-atom AMBER99DISP and CHARMM36m force-fields. The values corresponding to the all-atom force-fields are taken from Robustelli *et al.*[23]

| Force-field | $R_g$ (nm) | Reference |
|---|---|---|
| AMBER ff19SB/OPC | 1.84 | [24] |
| AMBER ff19SB/TIP4P-D | 3.13 | [24] |
| AMBER ff03CMAP/TIP4P-D | 2.03 | [24] |
| AMBER ff99DISP/TIP4P-D | 2.29 | [24] |
| AMBER ff12SB/TIP3P | 1.54, 1.90* | [25] |
| AMBER ff99SB-ILDN/TIP3P | 1.53, 1.60* | [25] |
| AMBER ff99SB-ILDN/TIP4P-EW | 1.79, 2.40* | [25] |
| AMBER ff99SB-ILDN/TIP4P-D | 2.57, 3.10* | [25] |
| AMBER ff12SB/TIP4P-D | 2.97, 3.41* | [25] |
| AMBER ff03ws/TIP4P | 3.0, 3.43* | [25] |
| AMBER ff99SBdisp/TIP4P-D | 2.62, 3.19* | [25] |
| CHARMM22*/TIP3P | 1.71, 2.31* | [25] |
| CHARMM22*/TIP4P-D | 2.33, 2.93* | [25] |
| SOP-IDP | 3.67 | This work |
| Bead-necklace | 3.60 | [26] |
| MARTINI | 2.88 | [26] |

Table S4: The $R_g$ values for $\alpha$-synuclein predicted by different force-fields. * denotes that the ensemble was reweighted prior to computing the $R_g$ values.

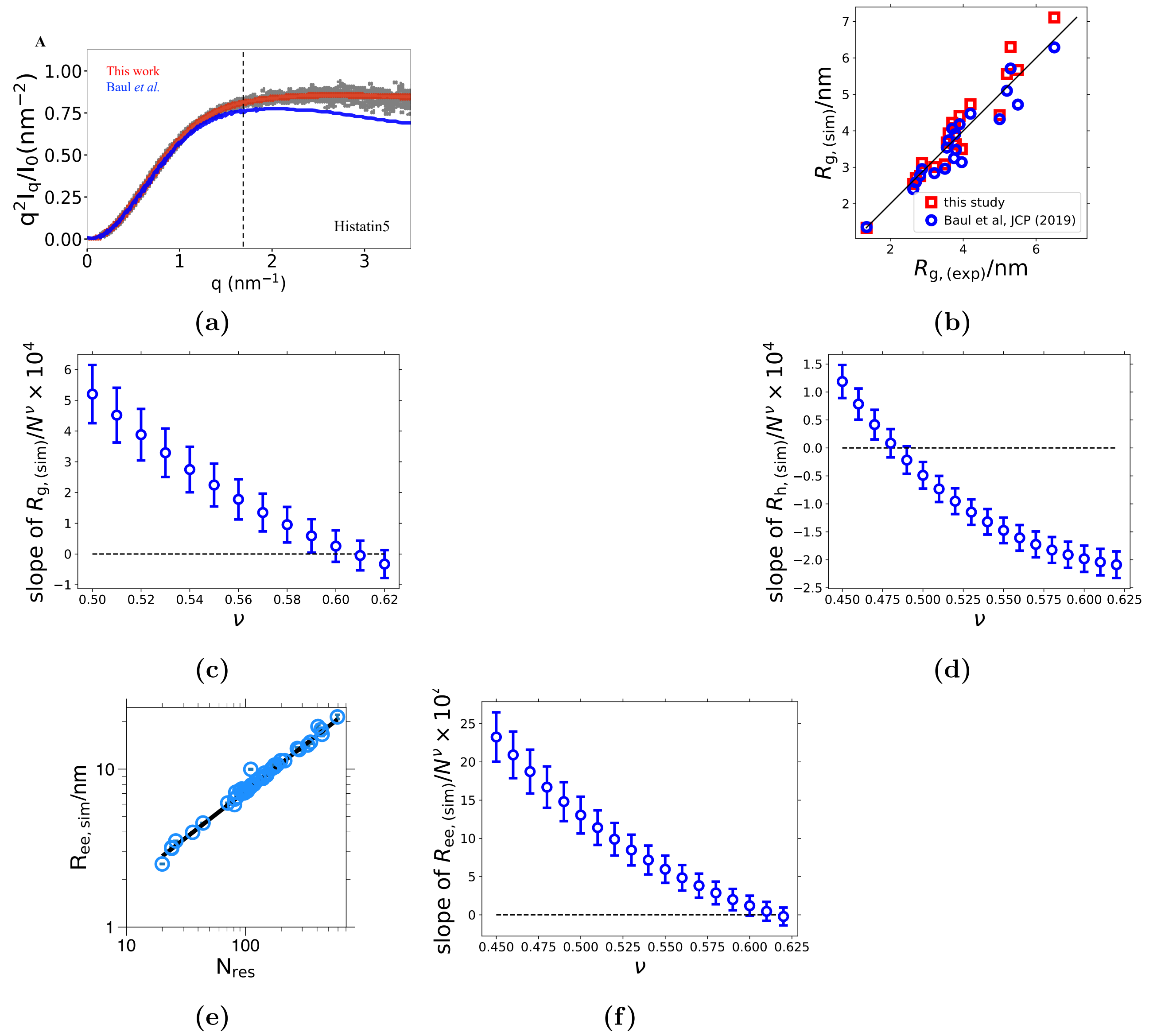


**Figure S1.** (A) Comparison between the SAXS profiles obtained with the current force-field (shown in red), and our previous work (shown in blue) for Histatin5. With the previous parameter set, [27] there are deviations from the experimental SAXS profile (grey points) beyond q $\approx$ 1.7 (shown by a dashed line). (B) Comparison between the experimental $R_g$ values and those obtained from previous simulations [27] (blue dots) and current model (red squares). The points represent 21 proteins used both in this and in the earlier study. The Pearson correlation coefficients are nearly identical, 0.95 and 0.96 for the earlier and new model. The solid green line provides a guide to eye. (C) Results of the fit of $R_g/N^{\nu}$ as a function of $N$ for various values of $\nu$. The y-axis shows the slope of this fit, which is expected to be $= 0$ (dashed black lines) for the correct exponent. (D) Same as (C), but for $R_h$. (E) End-to-end distance as a function of the number of residues. The fit performed with $\nu = 0.588$ gives a prefactor $a_e = (0.48 \pm 0.03)$ nm. (F) Same as (C), but for $R_{ee}$.

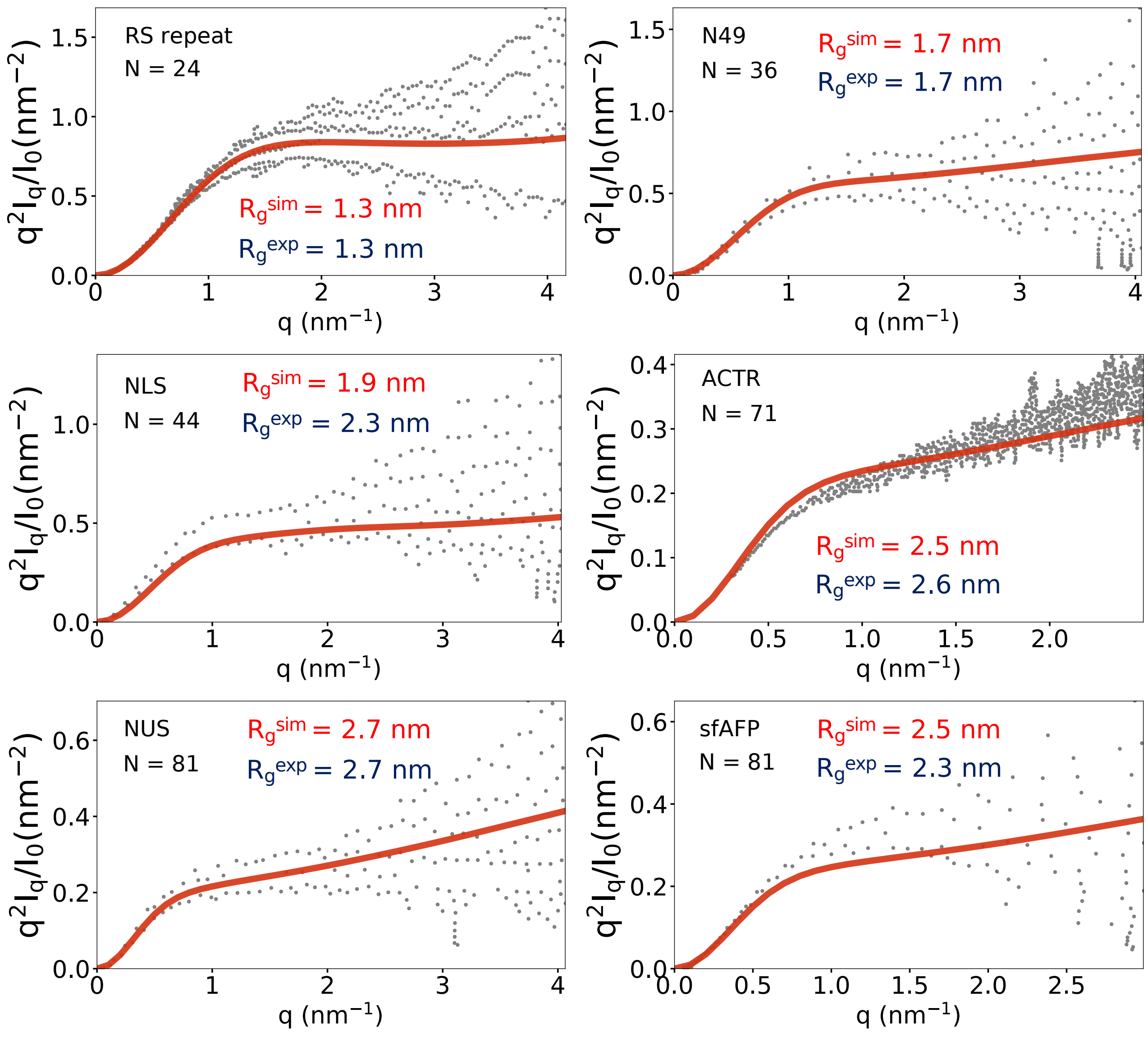


**Figure S2.** Kratky plots for RS repeat, N49, NLS, ACTR, NUS and sfAFP. The simulated profiles are shown in red, and the grey points denote the experimental data.

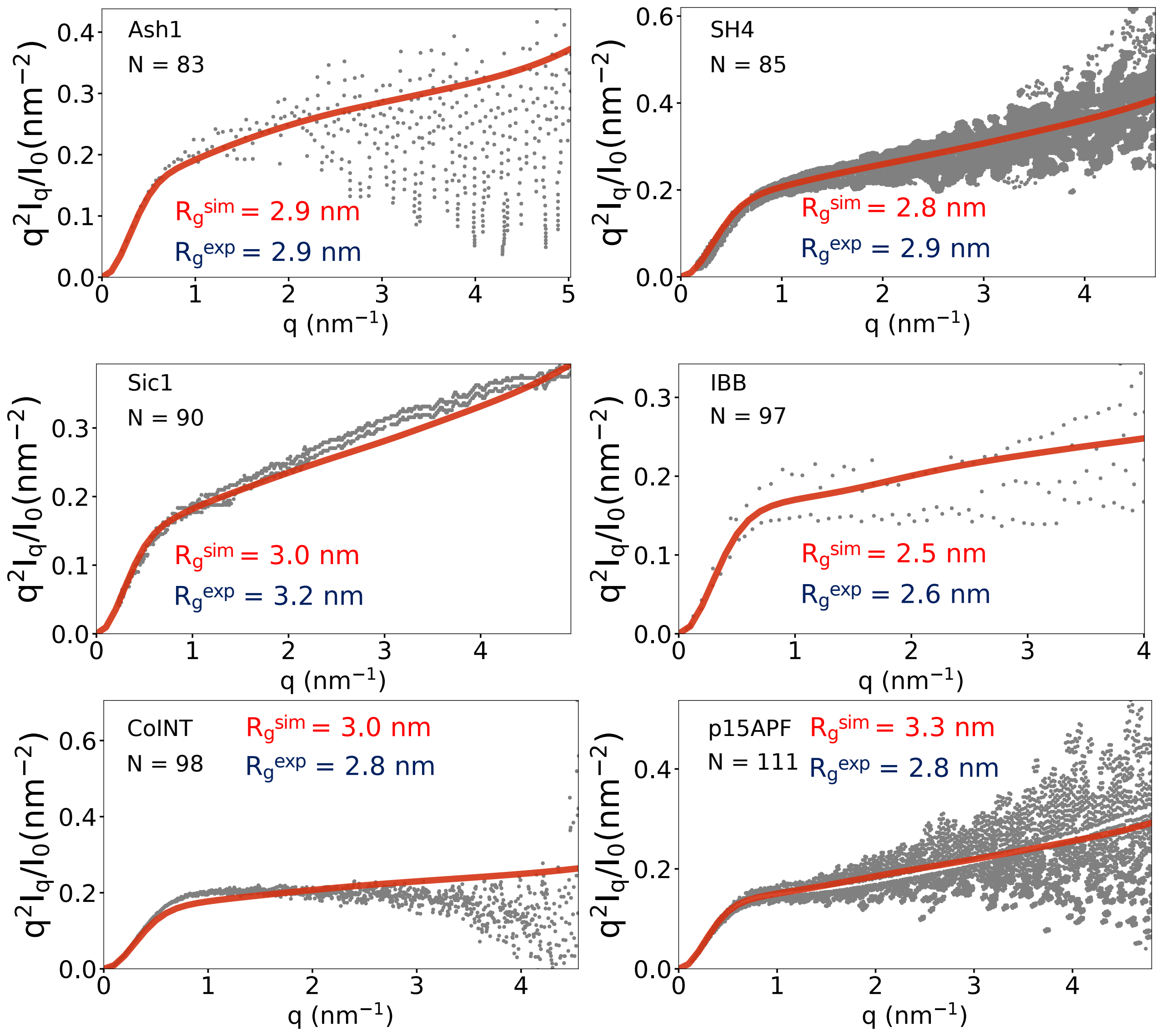


**Figure S3.** Kratky plots for Ash1, SH4, Sic1, IBB, ColNT and p15APF . The simulated profiles are in red, and the grey points denote the experimental data.

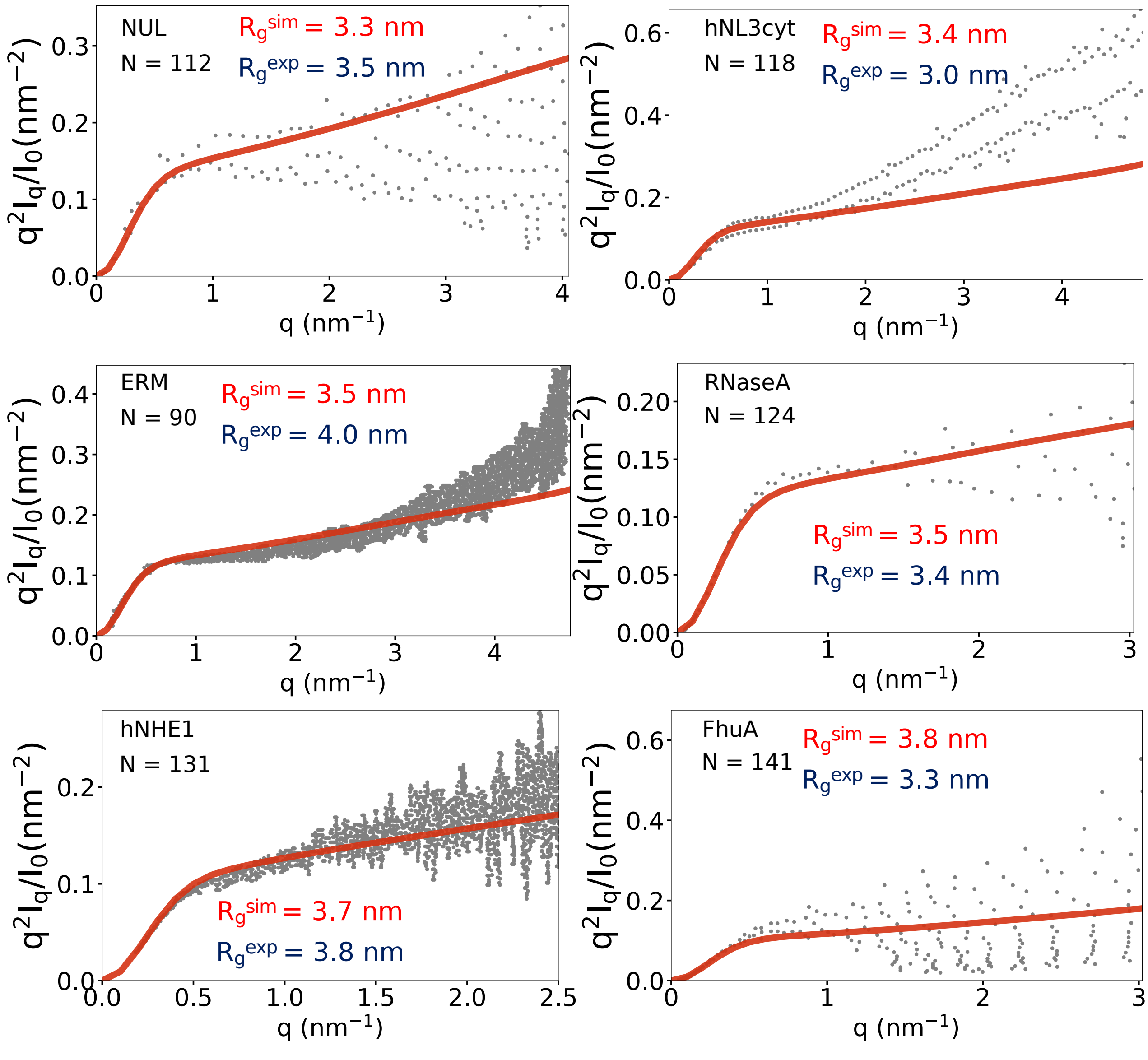


**Figure S4.** Kratky plots for NUL, hNL3cyt, ERM, RNaseA, hNHE1 and FhuA . The simulated profiles are shown in red, and the grey points denote the experimental data.

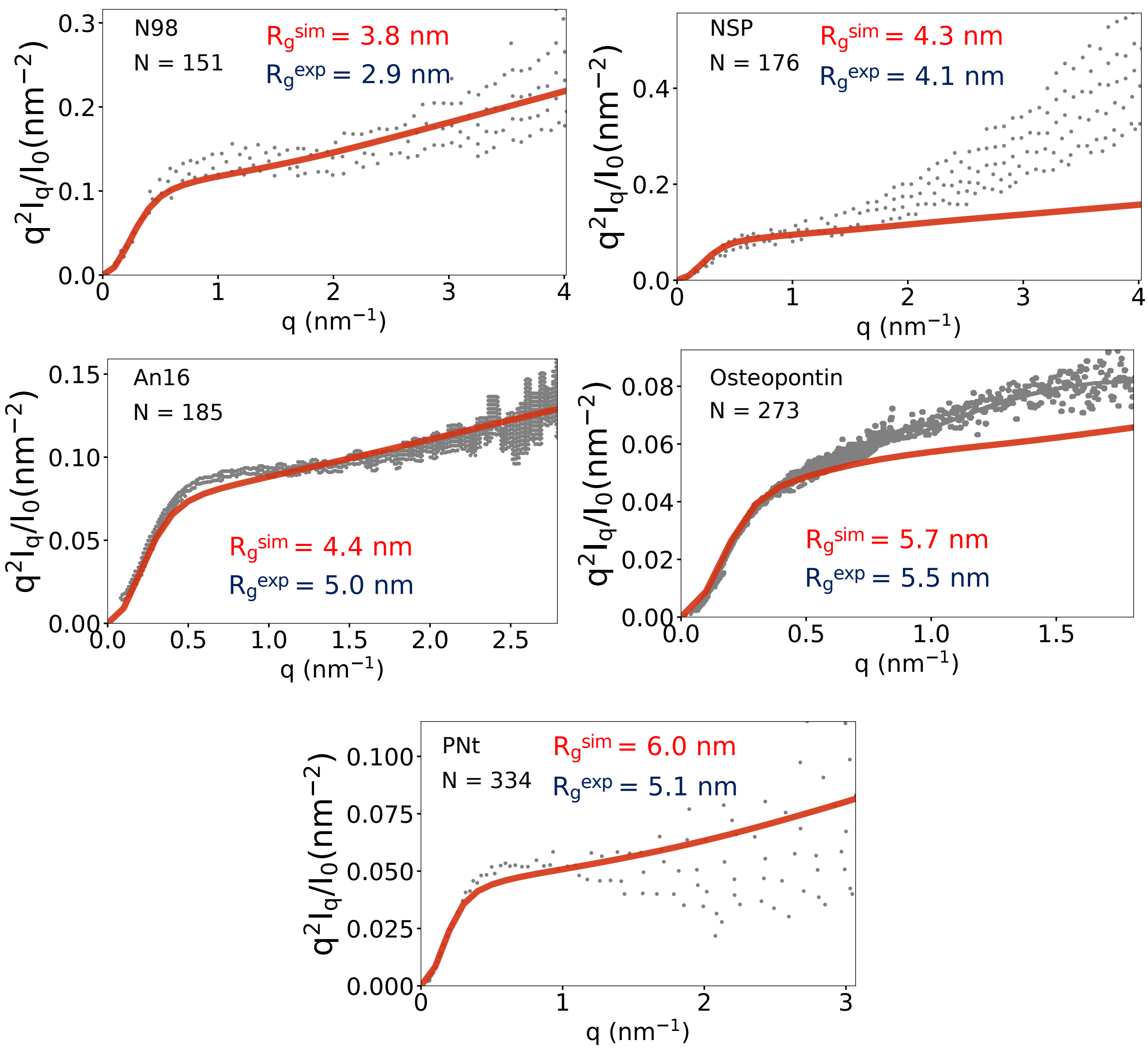


**Figure S5.** Kratky plots for N98, hNL3cyt, NSP, An16, Osteopontin and PNt . The simulated profiles are shown in red, and the grey points denote the experimental data.

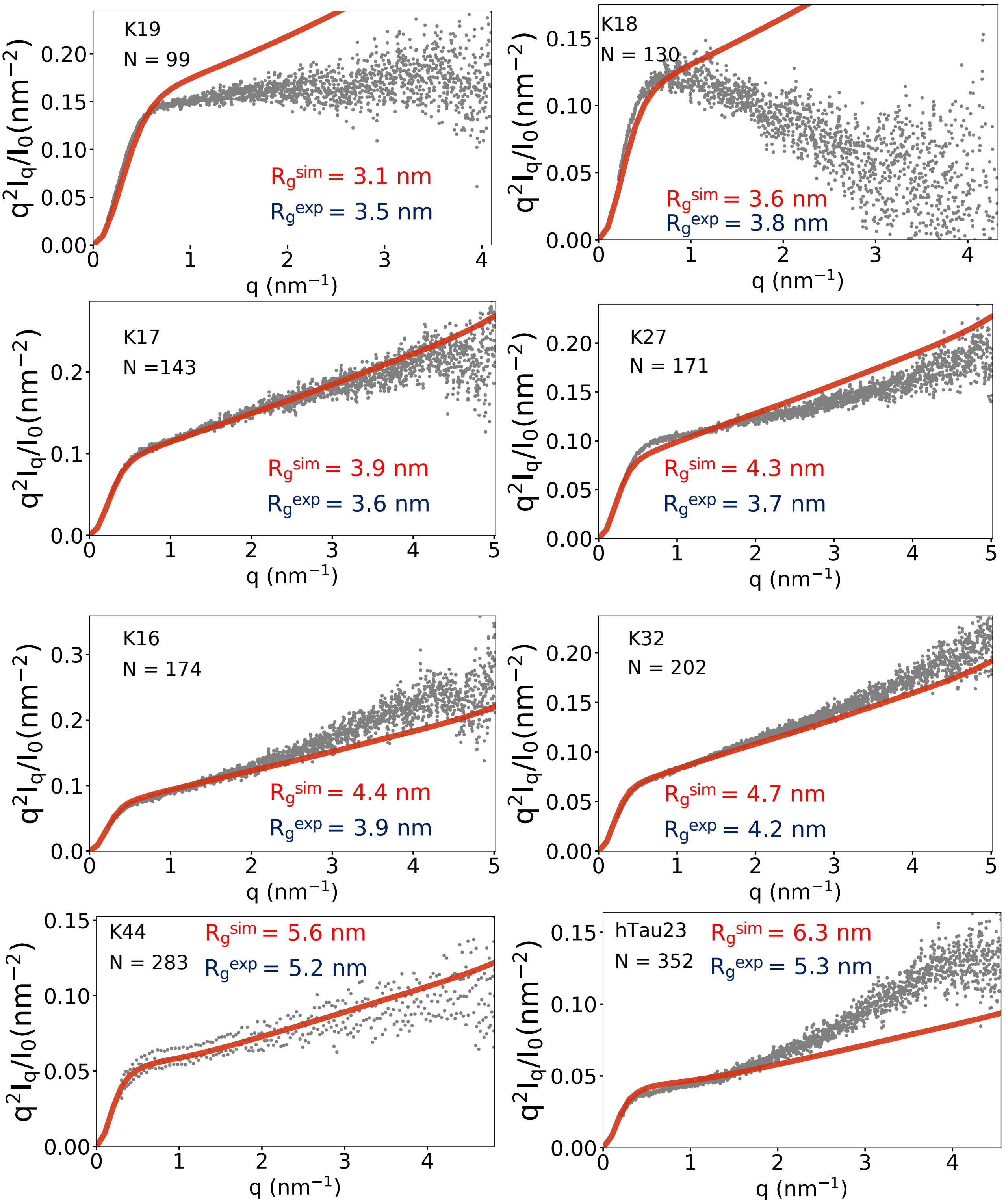


**Figure S6.** Kratky plots for the seven tau proteins, K19, K18, K17, K27, K16, K32, and hTau23 . The simulated profiles are shown in red, and the grey points denote the experimental data.

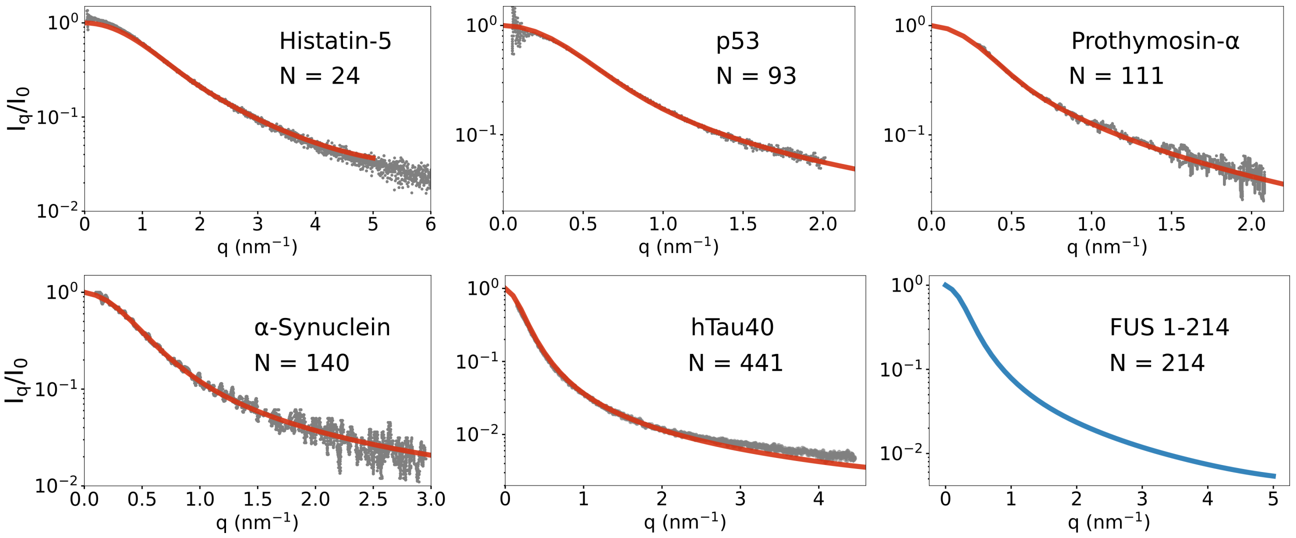


**Figure S7.** Scattering profiles for, Histatin-5, p53, Prothymosin-$\alpha$, $\alpha$-Synuclein, hTau40. The simulated profiles are shown in red, and grey points denote the experimental data. The SAXS profile of FUS 1-214 (shown in blue) serves as a prediction. Because the experimental SAXS curves are reported in arbitrary units, the simulated profiles are scaled by a constant factor to optimize the overlap with the experimental curves in the logarithmic scale.

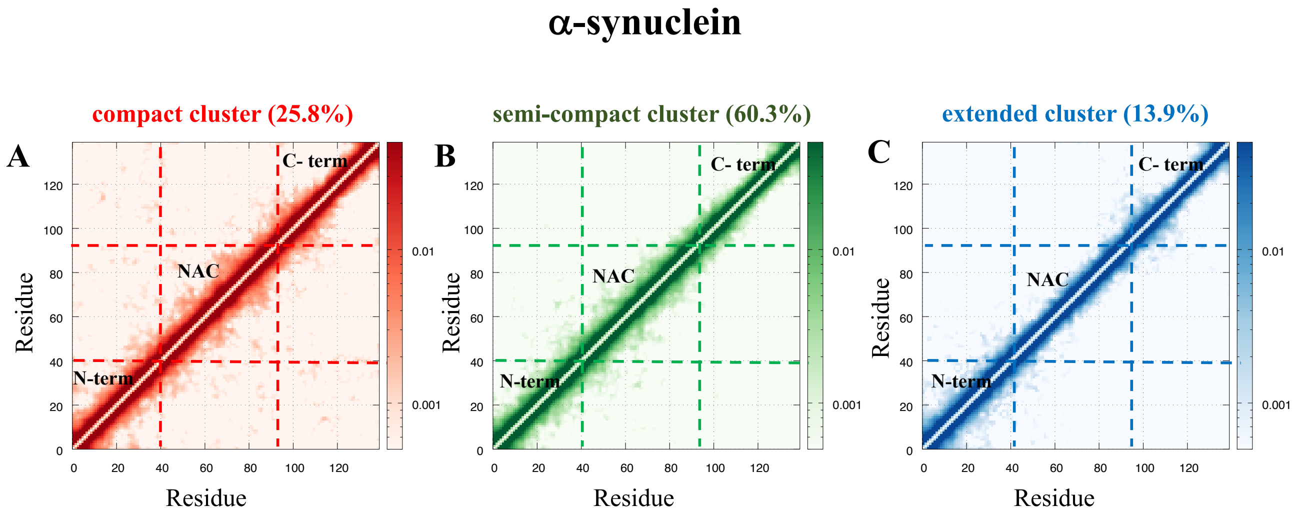


**Figure S8.** Difference contact maps for the (A) compact (population: 25.8%) (B) semi-compact (population: 60.3%) and (C) extended clusters (population: 13.9%) within the monomer conformational ensemble of $\alpha$-synuclein. The contact maps are color-coded according to the scheme in Figure 3, where red represents the most compact cluster, green represents the semi-compact clusters, and blue represents the extended cluster. The dashed lines on the contact maps demarcate the different segments of the $\alpha$-synuclein sequence, namely, the N-terminus (residues 1-40), the non-amyloid $\beta$-component (NAC) domain (residues 41-95), and the C-terminus (residues 96-140).

### htau40

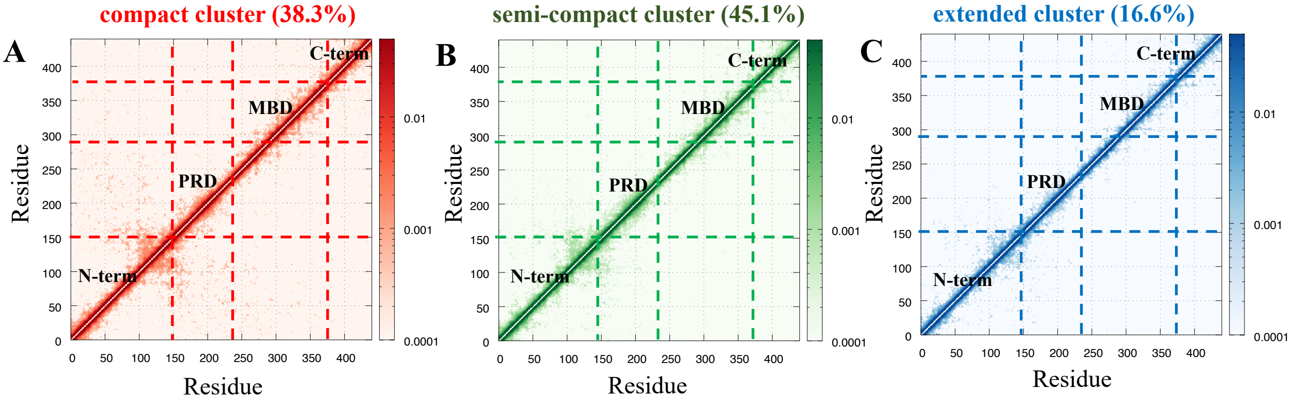


**Figure S9.** Difference contact maps for the (A) compact (population: 38.3%) (B) semi-compact (population: 45.1%) and (C) extended clusters (population: 16.6%) within the monomer conformational ensemble of htau40. The contact maps are color-coded according to the scheme in Figure 3, where red represents the most compact cluster, green represents the semi-compact clusters, and blue represents the extended cluster. The dashed lines on the contact maps demarcate the four segments of the htau40 sequence, namely, the N-terminus (residues 1-150), the proline-rich domain (PRD) (residues 151-244), the microtubule binding domain (MBD) (residues 245-372) and the C-terminus (residues 372-441).

# AP180

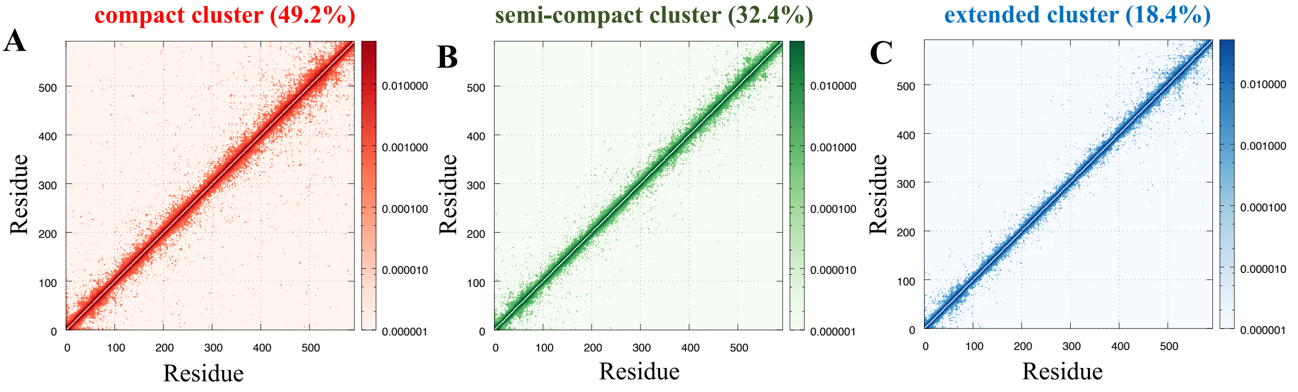


**Figure S10.** Difference contact maps for the (A) compact (population: 49.2%) (B) semi-compact (population: 32.4%) and (C) extended clusters (population: 18.4%) within the monomer conformational ensemble of AP180. The contact maps are color-coded according to the scheme in Figure 3, where red represents the most compact cluster, green represents the semi-compact clusters, and blue represents the extended cluster. Long-range contacts between the different segments of AP180 are not readily discernible from these contact maps.

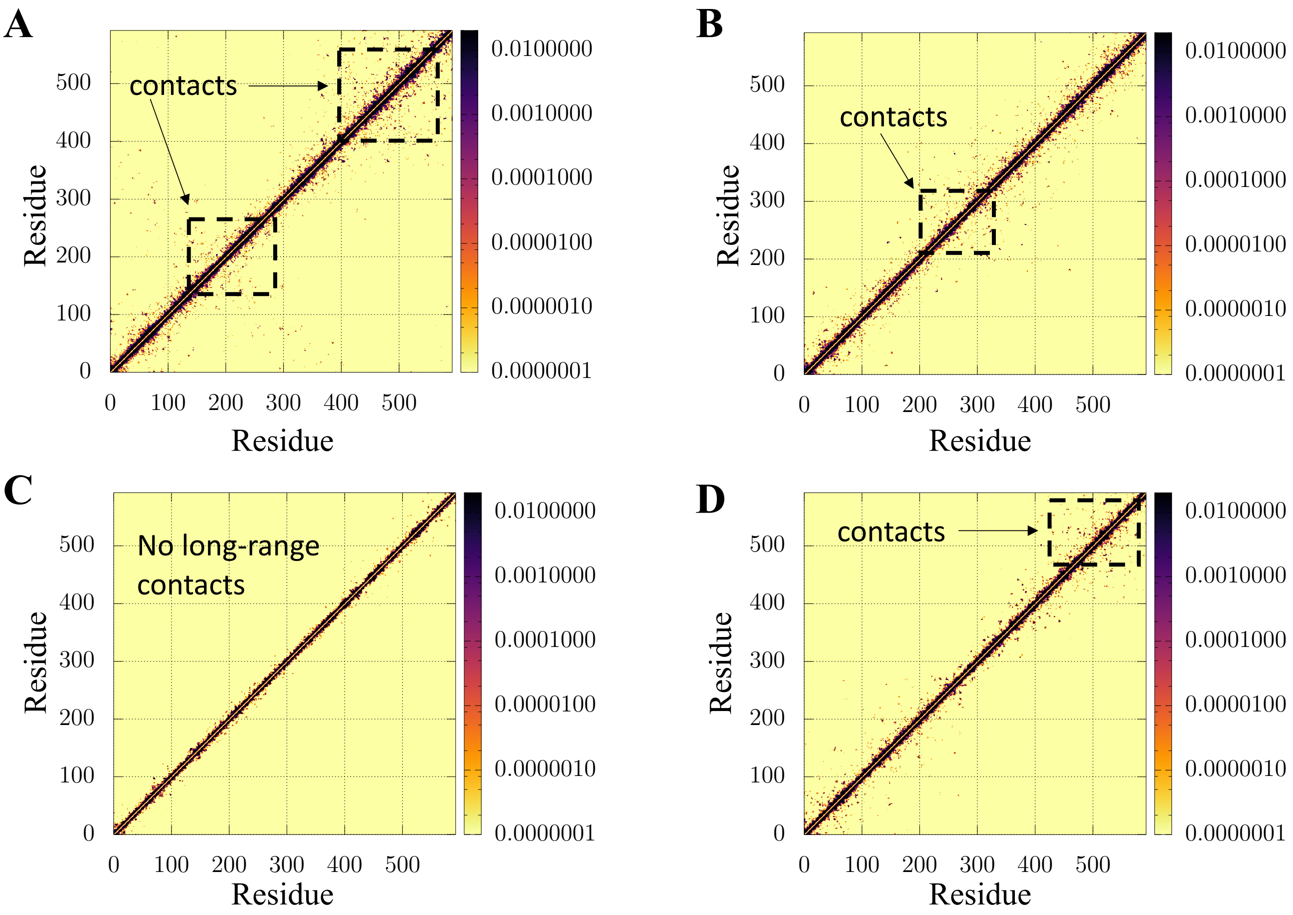


**Figure S11.** Contact maps corresponding to the different subclusters of AP180, obtained from a finer partitioning of the conformaitonal space. (A) A subfamily within the compact cluster exhibiting enhanced contacts at regions near the C-terminus, and the central segment. (B) A subfamily within the semi-compact cluster exhibiting enhanced contacts near the central segment. (C) A subfamily within the extended cluster having no long-range contacts, and exhibiting a contact map reminiscent of random-coils. (D) A subfamily within the extended cluster exhibiting contacts near the C-terminus.

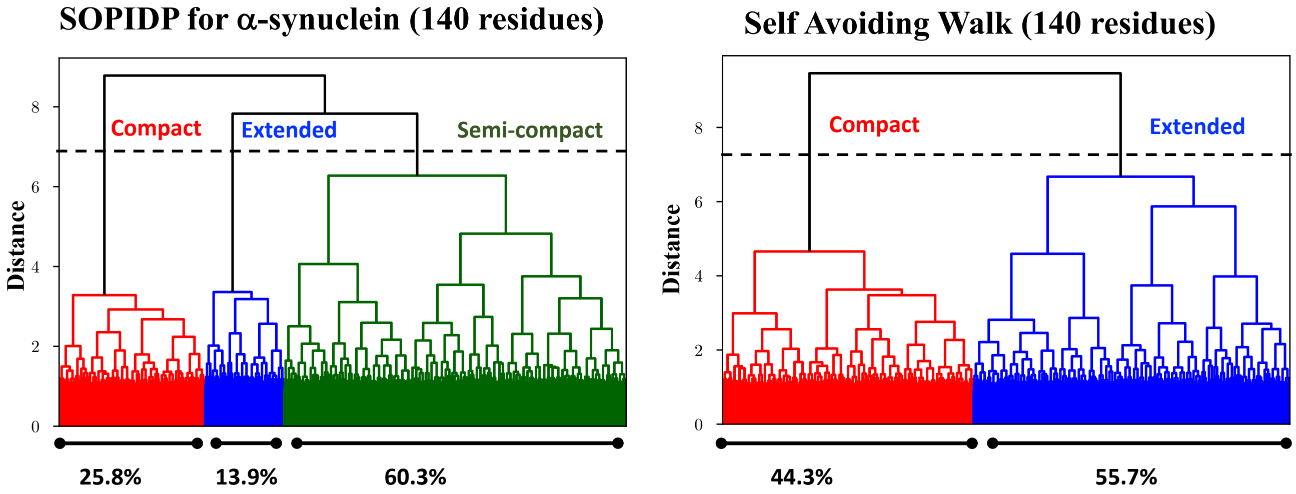


**Figure S12.** Hierarchical organization of clusters within the conformational ensembles of $\alpha$-synuclein (left) and a self-avoiding walk (SAW) (right) having 140 residues, depicted in the form of dendrograms. The dashed line denotes the distance cutoff used in both cases to segregate the different clusters. The SAW ensemble appears more homogeneous at the same distance cutoff, with a smaller difference in the populations of compact and extended clusters.
